# An analytical framework for phenotypic selection of fitness-conferring genes

**DOI:** 10.64898/2026.03.03.709325

**Authors:** Marc Sturrock, Anna Sturrock

**Affiliations:** Department of Physiology and Medical Physics, Royal College of Surgeons in Ireland, Dublin, Ireland

## Abstract

Phenotypic selection can cause the transient, selective upregulation of fitness-conferring genes in isogenic cell populations under stress, producing selective enrichment of the fitness gene relative to a neutral reference gene. While computational models have shown that such enrichment requires noisy gene expression and a cellular memory linking growth rate to gene expression (Ciechonska et al., 2022), the precise mechanistic requirements and the analytical principles governing enrichment have remained unclear. Here, we present an exact analytical framework that unifies enrichment mechanisms across both growth-driven and death-driven selection regimes. By analysing a stochastic model of explicit mRNA and protein dynamics, we prove that when selection acts via cell division, the fitness advantage of faster growth is exactly cancelled by the penalty of faster protein dilution. We show this symmetry is broken by translational feedback but not by transcriptional feedback alone; for genes with regulated (switching) promoters, the promoter-state memory provides an independent route to enrichment without translational feedback. Conversely, when selection acts via cell death, this exact cancellation is bypassed, allowing selective enrichment to emerge from baseline gene expression noise without any assumptions about growth-related feedback loops or regulated vs constitutive expression. We derive an exact fluctuation–response relation demonstrating that, in all cases, enrichment scales with the super-Poissonian component of unperturbed protein noise times the relevant memory timescale. All analytical predictions are corroborated by stochastic simulations of a finite-population Moran model. These results have implications for the emergence of drug resistance: by transiently enriching survival-conferring phenotypes, phenotypic selection can extend the window during which cell division occurs under stress, increasing the opportunity for permanent genetic mutations to arise.

## 1 Introduction

Genetically identical cell populations can exhibit phenotypic heterogeneity due to stochastic fluctuations in gene expression (Elowitz et al., 2002; Swain et al., 2002). Under environmental stress, this heterogeneity provides raw material for phenotypic selection: cells that happen to express higher levels of a fitness-conferring gene divide more frequently or survive longer, and thereby increase in relative abundance within the population. This leads to an apparent upregulation of the fitness gene at the population level, a phenomenon we term “selective enrichment” in this paper (Tsuru et al., 2011; Ciechonska et al., 2022).

Selective enrichment is distinct from genetic mutation, signal transduction, and bistable switching. It can operate even on a monostable, constitutively expressed gene, relying solely on the interplay between stochastic gene expression and fitness-dependent selection (via differential growth or survival). A natural way to quantify selective enrichment is to compare the fitness gene against a reference gene expressed with identical parameters in the same cell but whose protein does not affect the cell’s fitness. Since both genes have identical expression parameters and share the same intracellular environment, the ratio ⟨*n*⟩*/*⟨*r*⟩ of their mean protein levels isolates the effect of selection: any deviation from unity is a direct signature of selective enrichment. Understanding the mechanisms of selective enrichment is important because, although phenotypic selection is transient and non-genetic, it can extend the survival of subpopulations under stress, providing additional rounds of cell division during which permanent genetic resistance mutations may arise. This positions phenotypic selection as a potential stepping stone toward heritable drug resistance, both in bacterial antibiotic treatment and in chemotherapy applied to cancer cells (Ciechonska et al., 2022).

The analytical foundations for understanding phenotypic selection on gene expression were laid by Mora and Walczak (2013), who studied a birth–death model for protein copy number *n* with an *n*-dependent growth rate (denoted *s*(*n*) in their notation, not to be confused with the selection strength parameter *s*). For linear selection *s*(*n*) = *sn*, they showed that the steady-state distribution remains Poisson but with a shifted mean ⟨*n*⟩ = *b/*(*d* − *s*), where *b* is the protein production rate and *d* is the degradation rate. Compared to a reference gene unaffected by selection (⟨*r*⟩ = *b/d*), this gives a ratio ⟨*n*⟩*/*⟨*r*⟩ = *d/*(*d* − *s*) *>* 1 for positive selection, demonstrating selective enrichment.

However, the Mora–Walczak model treats protein dilution via a constant effective degradation rate *d*, independent of the cell’s growth rate. In reality, protein dilution occurs at cell division, when molecules are partitioned between daughter cells: a cell that grows faster divides sooner and dilutes its proteins more rapidly. This coupling between growth rate and dilution has important consequences for selective enrichment, as we demonstrate below (Huh and Paulsson, 2011; Thomas and Shahrezaei, 2021).

Computational agent-based models incorporating explicit cell division, stochastic gene expression, and various biological feedbacks were used to systematically investigate selective enrichment (Ciechonska et al., 2022). That study identified two key ingredients: variability in gene expression within an isogenic population, and a cellular “memory” from positive feedbacks between growth and expression. More specifically, their systematic comparison of ten candidate models revealed that, for growth-driven selection, enrichment required: (i) explicit modelling of both mRNA and protein (a two-stage model), (ii) growth-rate-dependent dilution, and (iii) a global positive feedback coupling the translation rate to the cell growth rate. Notably, a model with a growth-rate-dependent transcription rate but a growth-rate-independent per-mRNA translation rate (Model 6 of Ciechonska et al. 2022) did not produce selective enrichment, and Bayesian model comparison assigned the model with growth-dependent translation alone (Model 7) as most likely to produce enrichment, as opposed, perhaps counter-intuitively, to the model with both transcriptional and translational feedbacks (Model 8).

In this paper, we provide an analytical theory that explains these computational findings. We begin by proving two no-enrichment theorems for growth-coupled dilution (Theorems 1 and 2), demonstrating that when protein dilution is coupled to the growth rate but there is no translational feedback, the ratio ⟨*n*⟩*/*⟨*r*⟩ = 1 exactly, for any selection strength *s* ≥ 0. This holds regardless of whether transcriptional feedback is present, and requires no moment closure approximation. We then extend the model to include explicit mRNA dynamics and a global translational feedback (translation rate scales with growth rate), showing that selective enrichment emerges. For a general model with transcriptional feedback *q*_1_ and translational feedback *q*_2_, the enrichment ratio at weak selection is

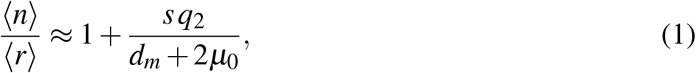

which is independent of *q*_1_ at leading order. To generalise this result, we derive an exact (all-orders) relation showing that selective enrichment is proportional to *q*_2_ times the covariance asymmetry Cov(*n, m*_*n*_) −Cov(*n, m*_*r*_). The underlying mechanism can be understood in terms of timescales: translational feedback exploits the mRNA autocorrelation time *τ*_*m*_ = 1*/*(*d*_*m*_+*µ*_0_) to amplify gene-specific stochastic memory. The intrinsic covariance *C*_*nm*_ = Cov(*n, m*_*n*_) between a protein and its own mRNA (Thattai and van Oudenaarden, 2001; Paulsson, 2005) provides the memory window; translational feedback converts this memory into a selective advantage because it multiplies the gene’s own mRNA (*m*_*n*_ or *m*_*r*_). Transcriptional feedback, by contrast, acts upstream of the gene-specific mRNA and creates no gene-specific memory that persists across cell division events. We further show that when selection acts via differential cell death rather than differential growth, this exact cancellation is bypassed entirely, and selective enrichment emerges from baseline gene expression noise without any translational feedback. All analytical predictions are corroborated by stochastic simulations of a finite-population Moran model.

## 2 Model framework

We work within the population-level framework of Mora and Walczak (2013). A population of cells is described by the density *ρ*_**x**_(*t*), where **x** denotes the intracellular state of a cell. Within each cell, biochemical reactions govern the stochastic dynamics of gene expression.

To keep the population size constant (as in a chemostat or turbidostat; Gresham and Dunham 2014), each cell division displaces a randomly chosen cell, creating an effective death rate equal to the population-mean growth rate ⟨*µ*⟩. The normalised probability *P*_**x**_(*t*) = *ρ*_**x**_(*t*)*/* ∑_**x**_′ *ρ*_**x**_′ (*t*) then satisfies

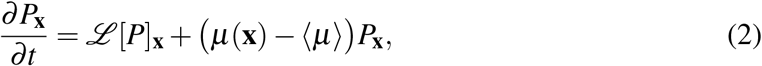

where ℒ is the operator describing intracellular birth–death dynamics.

Each cell contains two genes: a fitness gene producing protein *n* and a reference gene producing protein *r*. Both genes are constitutively expressed with identical parameters. The cell’s growth rate depends only on the fitness protein: *µ*(*n*) = *µ*_0_+*sn*, where *µ*_0_ is the basal growth rate and *s >* 0 quantifies the selection strength. In Section 3.6, we generalise this framework to include death-driven selection, where the fitness protein reduces a cell’s death rate rather than increasing its growth rate.

In Models A–E below, both genes are constitutively expressed (no promoter regulation); we relax this assumption in Section 3.5. We consider five models of increasing biological realism.

### 2.1 Model A: Mora–Walczak baseline (protein only, constant dilution)

Protein dynamics follow a simple birth–death process with constant rates. There is no explicit mRNA. Proteins are produced at a constant rate *b* and degraded at rate *d* per molecule, giving a total degradation rate *d* · *n* per cell. The growth rate is *µ*(*n*) = *sn*. This is the model studied by Mora and Walczak (2013). Dilution is absorbed into *d*, independent of growth rate.

### 2.2 Model B: Growth-coupled dilution (protein only, no feedback)

The per-molecule dilution rate equals the cell’s growth rate. There is no explicit mRNA. Proteins are produced at a constant rate *b* and diluted at rate *µ*(*n*) per molecule, giving a total dilution rate *µ*(*n*) · *n* per cell. The growth rate is *µ*(*n*) = *µ*_0_+*sn*.

### 2.3 Models C–E: Growth-coupled dilution with explicit mRNA and feedback

We now include explicit mRNA dynamics and allow for two types of global positive feedback: growth-dependent transcription (strength *q*_1_) and growth-dependent translation (strength *q*_2_). These are motivated by evidence that both RNA polymerase abundance and ribosome abundance scale with growth rate in bacteria (Klumpp and Hwa, 2008; Klumpp et al., 2009; Dai et al., 2016; Shahrezaei and Marguerat, 2015).

For each gene (*n* or *r*), the intracellular reactions with growth rate *µ*(*n*) = *µ*_0_+*sn* are:

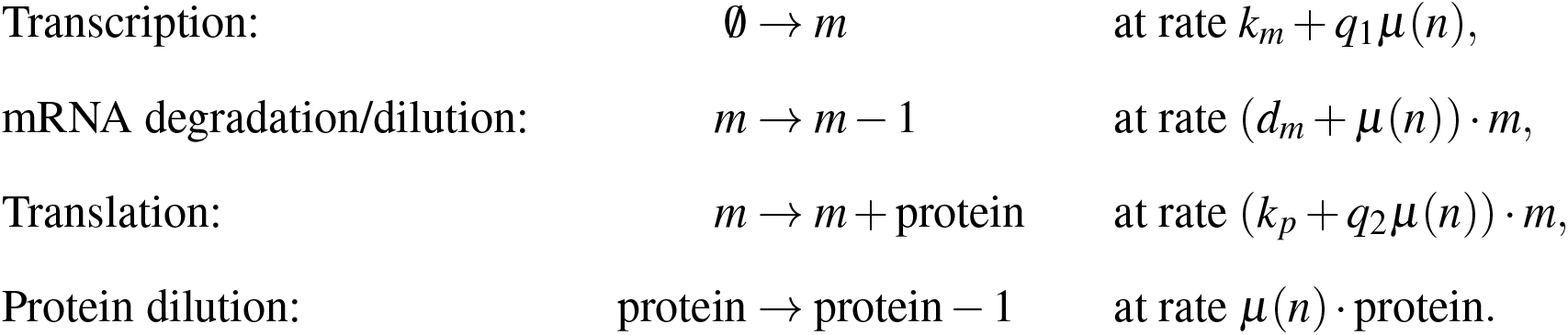

Three special cases are of interest, corresponding directly to Models 6–8 of Ciechonska et al. (2022): Model C (transcriptional feedback only, *q*_1_ *>* 0, *q*_2_ = 0), Model D (translational feedback only, *q*_1_ = 0, *q*_2_ *>* 0), and Model E (both feedbacks, *q*_1_ *>* 0, *q*_2_ *>* 0).

### 2.4 General moment equation

For any intracellular quantity *ϕ* (**x**), the evolution of its population mean under Eq. (2) is

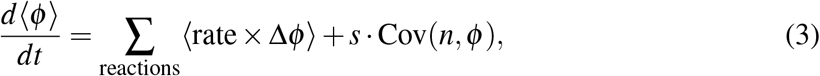

where Δ*ϕ* is the change in *ϕ* due to each reaction. With linear selection *µ*(*n*) = *µ*_0_+*sn*, the selection contribution takes the form *s* Cov(*n, ϕ*) = *s*(⟨*nϕ*⟩ −⟨*n*⟩⟨*ϕ*⟩), a manifestation of Fisher’s fundamental theorem (Fisher, 1930).

### 2.5 Model A: The Mora–Walczak result

Applying Eq. (3) to *ϕ* = *n*:

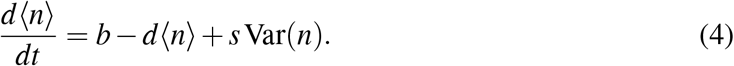

Since the steady-state distribution is Poisson (Var(*n*) = ⟨*n*⟩) (Mora and Walczak, 2013), the steady state gives

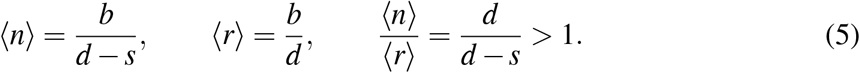

This is the main result of Mora and Walczak (2013): selection effectively reduces the degradation rate from *d* to *d* − *s*, producing selective enrichment.

## 3 Results

### 3.1 Growth-coupled dilution exactly cancels selection

#### Theorem 1

(No enrichment under growth-coupled dilution). *Consider a protein produced at constant rate b and diluted at the growth rate µ*(*n*), *where µ is any function of the fitness protein copy number n. Let the reference protein r be produced at the same rate b and diluted at the same growth rate µ*(*n*). *Then* ⟨*n*⟩*/*⟨*r*⟩ = 1 *at steady state, for any fitness landscape µ*(*n*).

*Proof*. We first prove the result for linear selection *µ*(*n*) = *µ*_0_+*sn*, then generalise. Applying Eq. (3):

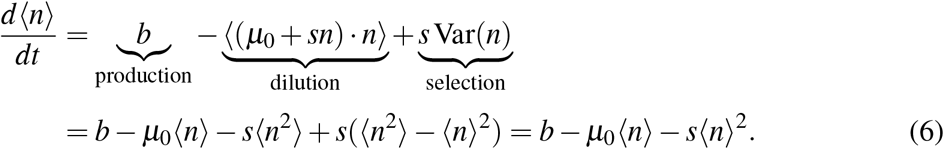

The ⟨*n*^2^⟩ terms from dilution and selection cancel exactly, leaving a closed equation for ⟨*n*⟩ that requires no moment closure. The steady state satisfies

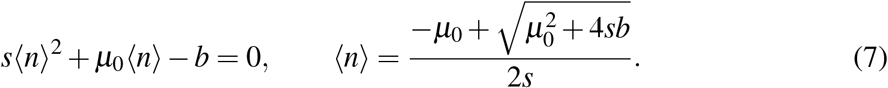

For the reference protein, the same cancellation yields

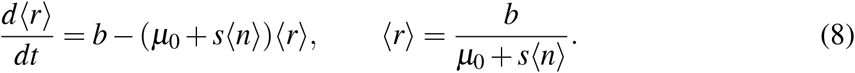

Using *b* = ⟨*n*⟩(*µ*_0_+*s*⟨*n*⟩) from Eq. (7):

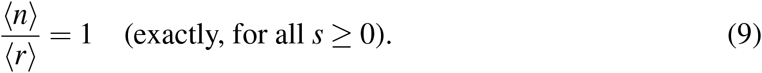

The cancellation does not rely on the linear form *µ*(*n*) = *µ*_0_+*sn*. For a general fitness landscape *µ*(*n*), the general moment equation (Appendix A) gives

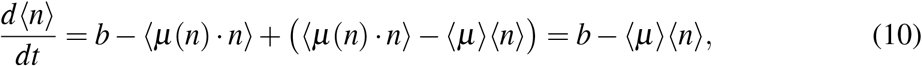

where the dilution term −⟨*µ*(*n*) · *n*⟩ and the selection term +⟨*µ*(*n*) · *n*⟩ − ⟨*µ*⟩⟨*n*⟩ annihilate the ⟨*µ*(*n*) · *n*⟩ dependence exactly for any *µ*(*n*). An identical argument gives *d*⟨*r*⟩*/dt* = *b* −⟨*µ*⟩⟨*r*⟩. Therefore ⟨*n*⟩*/*⟨*r*⟩ = 1 for any nonlinear fitness landscape.

Growth-coupled dilution completely eliminates selective enrichment: the selective advantage of faster-growing cells is exactly offset by their faster protein dilution. Figure 1 provides a distributional view of these results, showing how the protein copy number distributions for the fitness and reference genes differ across models; the top-right panel (Model B) shows the perfect overlap predicted by Theorem 1.

**Figure 1:**
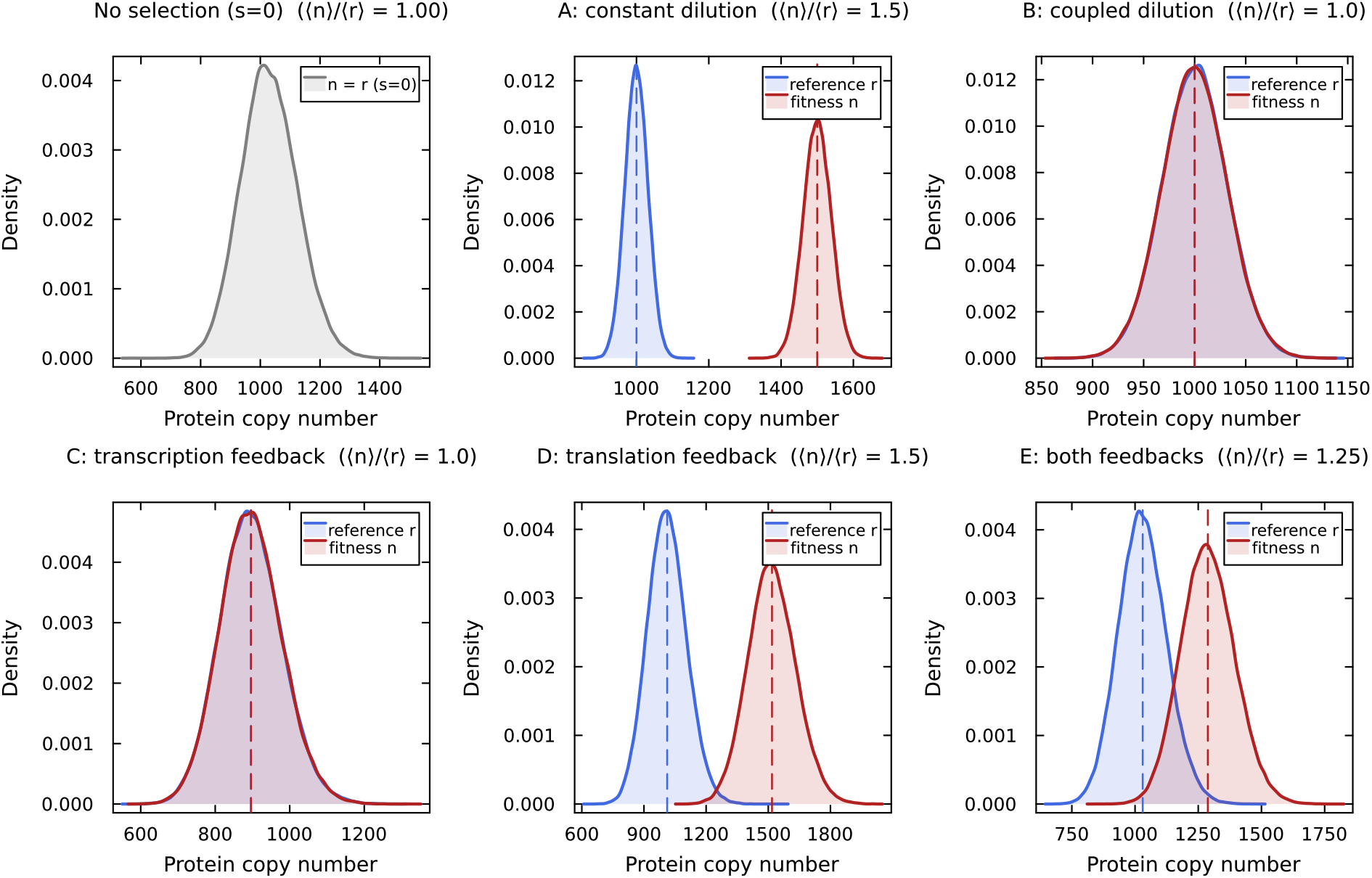
Distributional view of selective enrichment across models at selection strength *s* = 0.5. Each panel shows kernel density estimates of the protein copy number distribution for the fitness gene (red) and reference gene (blue), with dashed vertical lines at the means and the enrichment ratio annotated. Distributions are moment-matched analytical distributions (negative binomials for two-stage models C–E, Poissons for protein-only models A, B) with means from the Gaussian moment closure (Appendix D) for Models D and E. Model A (constant dilution) produces a clear rightward shift; Models B and C (growth-coupled dilution, with or without transcriptional feedback) show perfect overlap (⟨*n*⟩*/*⟨*r*⟩ = 1); Models D and E (translational feedback) restore the shift. The top-left panel shows the baseline distribution at *s* = 0.

### 3.2 Translational feedback is necessary and sufficient for selective enrichment

We now analyse the general model with mRNA and both feedbacks (*q*_1_, *q*_2_).

#### 3.2.1 mRNA means

Applying Eq. (3) to *ϕ* = *m*_*n*_:

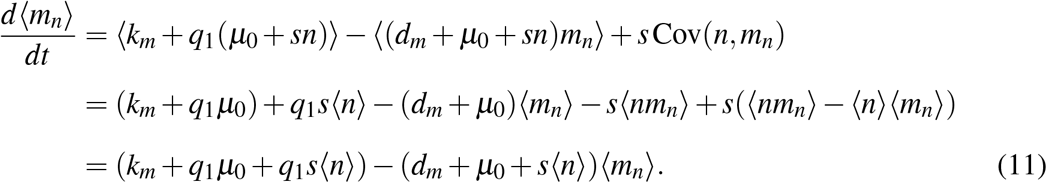

The raw cross-moment ⟨*nm*_*n*_⟩ cancels exactly. Note that the transcription feedback appears only through ⟨*n*⟩, which is the same for both genes. Therefore an identical equation holds for ⟨*m*_*r*_⟩, and at steady state:

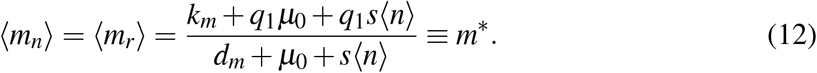

The mRNA means are always equal for the two genes, regardless of the transcriptional feedback strength *q*_1_.

#### 3.2.2 Protein means

For the fitness protein *n*, with translation rate (*k*_*p*_+*q*_2_(*µ*_0_+*sn*)) · *m*_*n*_:

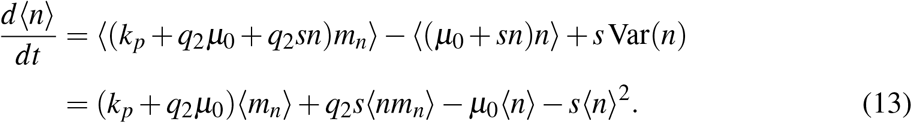

For the reference protein *r*:

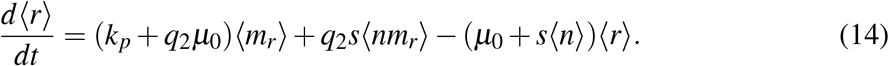

The key structural difference lies in the cross-moment terms. For the fitness protein, the term *q*_2_*s*⟨*nm*_*n*_⟩ involves *n* and its own mRNA *m*_*n*_, whereas for the reference protein, the term *q*_2_*s*⟨*nm*_*r*_⟩ involves *n* and the reference mRNA *m*_*r*_. These terms are present if and only if *q*_2_≠ 0. When *q*_2_ = 0 (no translational feedback), both protein equations become closed (requiring no higher-order moments) and reduce to the form of Model B.

#### 3.2.3 Model C: Transcriptional feedback only (*q*_1_ *>* 0, *q*_2_ = 0)

##### Theorem 2

(No enrichment under transcriptional feedback). *Consider the two-stage model with growth-coupled dilution and an arbitrary transcription function h*(*µ*) *of the growth rate, but with a growth-rate-independent translation rate (q*_2_ = 0*). Then* ⟨*n*⟩*/*⟨*r*⟩ = 1 *at steady state, for any fitness landscape µ*(*n*) *and any transcription function h*.

*Proof*. Setting *q*_2_ = 0, the protein equations become:

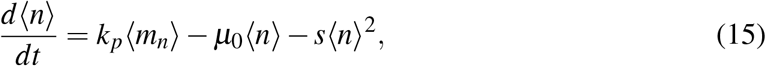

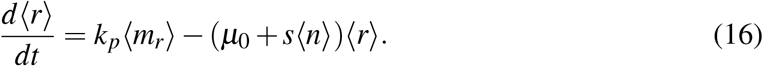

Since ⟨*m*_*n*_⟩ = ⟨*m*_*r*_⟩, these have identical source terms. By the same algebraic argument as Theorem 1:

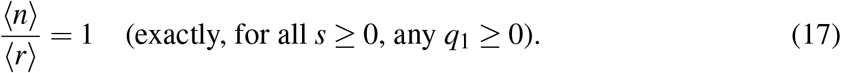

The transcription rate (*k*_*m*_+*q*_1_*µ*(*n*)) depends on *n* through the growth rate, but it does not multiply any gene-specific stochastic variable. The production of mRNA for gene *n* and gene *r* is affected identically by the growth-rate-dependent transcription enhancement, so the mRNA means remain equal and the downstream protein dynamics inherit this symmetry. For a general transcription function *h*(*µ*), the mRNA source ⟨*h*(*µ*(*n*))⟩ is always gene-independent, ensuring ⟨*m*_*n*_⟩ = ⟨*m*_*r*_⟩ and hence ⟨*n*⟩*/*⟨*r*⟩ = 1 regardless of the form of *µ* or *h*.

This result explains why Model 6 of Ciechonska et al. (2022), which had growth-dependent transcription but no translational feedback, produced a max ratio of exactly 1.0 in computational simulations. The bottom-left panel of Figure 1 (Model C) illustrates this: the fitness and reference distributions overlap perfectly despite the presence of transcriptional feedback.

#### 3.2.4 Models D and E: Translational feedback (*q*_2_ *>* 0)

When *q*_2_ *>* 0, the protein equations contain the unclosed cross-moments ⟨*nm*_*n*_⟩ and ⟨*nm*_*r*_⟩. We proceed by perturbative expansion in *s* (weak selection).

#### 3.2.5 Zeroth order (*s* = 0)

At zero selection, the growth rate is constant (*µ*_0_) and both genes are independent and identically distributed. The effective rates are:

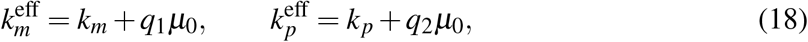

and the steady-state values are:

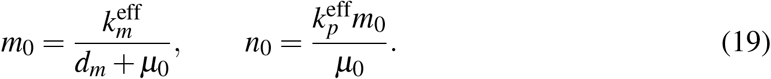

#### 3.2.6 Intrinsic protein–mRNA covariance

The covariance between a protein and its own mRNA at *s* = 0, computed from the Lyapunov equation (Appendix B), is

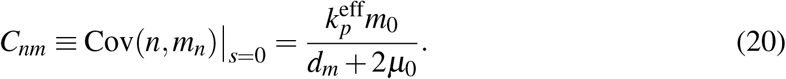

This is always positive: cells with more mRNA of a gene tend to have more of that gene’s protein. Importantly, the fitness protein *n* and the reference mRNA *m*_*r*_ are independent at *s* = 0:

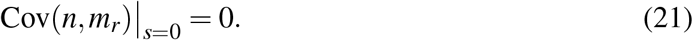

#### 3.2.7 First-order corrections

Expanding ⟨*n*⟩ = *n*_0_+*sn*_1_, ⟨*r*⟩ = *n*_0_+*sr*_1_, and similarly for mRNA means, the first-order steadystate conditions give (see Appendix C for details). For the fitness protein, the *O*(*s*) balance of Eq. (13) is

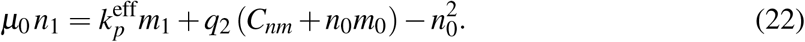

For the reference protein, using Cov(*n, m*_*r*_)|_*s*=0_ = 0:

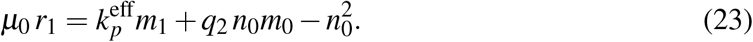

The terms 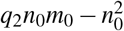 are common to both equations and cancel in the difference. The transcriptional feedback *q*_1_ enters only through *m*_0_ and *m*_1_, which are identical for both genes.

#### 3.2.8 The enrichment ratio

Subtracting Eqs. (22) and (23):

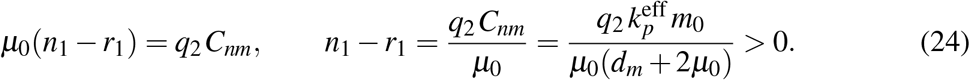

Dividing by 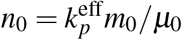, the ratio of means is

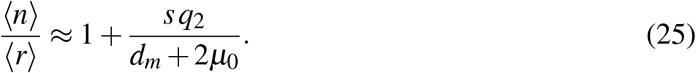

This is our main finding for the growth-dependent enrichment analysis. At leading order in *s*, the enrichment ratio is completely independent of the transcriptional feedback strength *q*_1_; only *q*_2_ (translational feedback) appears. The ratio reduces to unity when *q*_2_ = 0, consistent with the exact results for Models B and C. The Fano factor of the two-stage model at *s* = 0 is 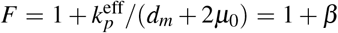, where 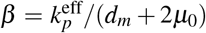 is the effective translational burst contribution (the ratio of the effective translation rate to the total mRNA turnover rate, incorporating growth-rate corrections to both). Writing 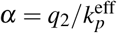, we have *q*_2_*/*(*d*_*m*_ + 2*µ*_0_) = *αβ*, and the enrichment ratio equals 1 + *sα*(*F* − 1): selective enrichment scales directly with the super-Poissonian component of protein noise.

#### 3.2.9 Exact relation at finite selection

Although Models D and E give identical enrichment ratios at first order in *s*, an exact (all-orders) relation reveals their structural difference. Subtracting the steady-state equations (13) and (14), and using ⟨*m*_*n*_⟩ = ⟨*m*_*r*_⟩ = *m*^∗^, yields

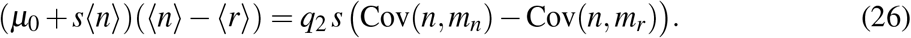

This relation holds exactly, without perturbative expansion or moment closure. It shows that the enrichment difference ⟨*n*⟩ − ⟨*r*⟩ is proportional to *q*_2_ at all orders in *s*, so that if *q*_2_ = 0 the right-hand side vanishes and Theorems 1 and 2 are recovered immediately. The driving force is the covariance asymmetry Δ*C* ≡ Cov(*n, m*_*n*_) −Cov(*n, m*_*r*_), which measures how much more strongly the fitness protein correlates with its own mRNA than with the reference mRNA. At *s* = 0, Cov(*n, m*_*n*_) = *C*_*nm*_ and Cov(*n, m*_*r*_) = 0, so Δ*C* = *C*_*nm*_ and the first-order result (25) is recovered. At finite *s*, both covariances shift; the effect of *q*_1_ on these covariance shifts is discussed below in the context of the signal-to-penalty decomposition.

#### 3.2.10 Second-order correction: transcriptional feedback suppresses selective enrichment

Although *q*_1_ is absent at first order, the exact relation (26) reveals how it enters at the next order. Rewriting Eq. (26) as

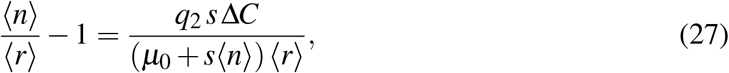

and approximating Δ*C* ≈ *C*_*nm*_ (its value at *s* = 0, valid at leading order), ⟨*n*⟩ ≈ ⟨*r*⟩ ≈ *n*_0_, gives

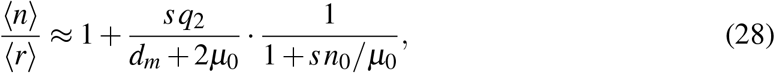

where the zeroth-order protein level

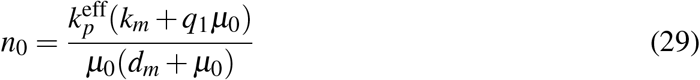

grows linearly with the transcriptional feedback strength *q*_1_. The factor 1*/*(1 + *sn*_0_*/µ*_0_) represents a nonlinear dilution penalty: at finite selection, the growth rate *µ*_0_+*s*⟨*n*⟩ exceeds the basal rate *µ*_0_, accelerating dilution. Transcriptional feedback inflates *n*_0_ (by boosting mRNA production via *k*_*m*_+*q*_1_*µ*_0_), which amplifies this penalty without proportionally strengthening the gene-specific signal Δ*C* in the numerator. The cancellation of *q*_1_ at first order occurs because *C*_*nm*_ ∝ *n*_0_ (both scale with *k*_*m*_+*q*_1_*µ*_0_), so the ratio *C*_*nm*_*/n*_0_ is *q*_1_-independent. At second order, however, the denominator grows as 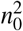 while the numerator grows only as *n*_0_, and the suppression becomes visible.

Expanding Eq. (28) to second order in *s*:

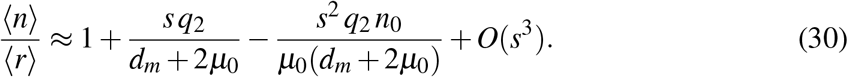

The second-order correction is strictly negative and its magnitude increases with *q*_1_ through *n*_0_. This provides the analytical basis for the suppression of selective enrichment by transcriptional feedback observed in the Gaussian moment closure and Moran simulations (Figure 2).

**Figure 2:**
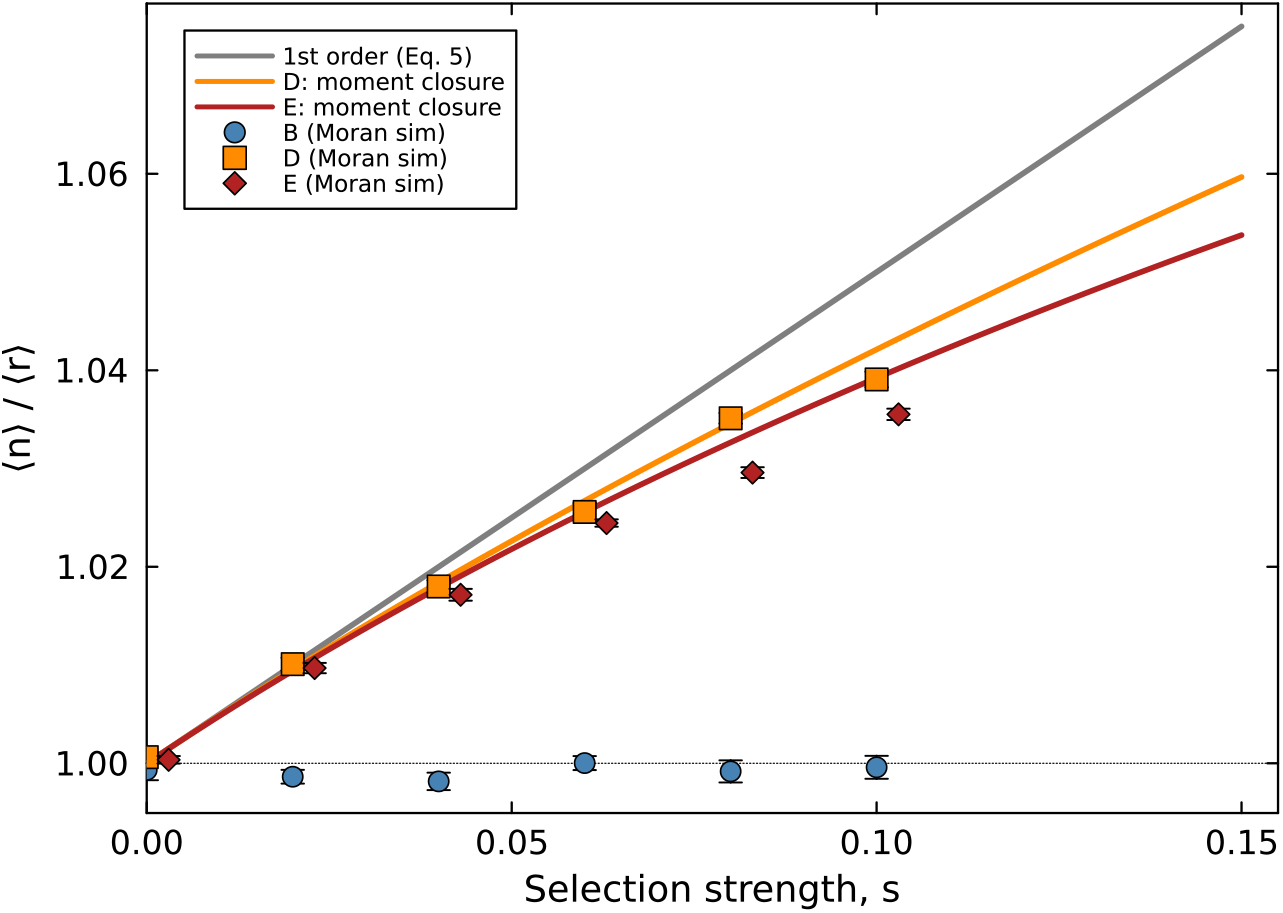
Validation of analytical predictions against Moran model simulations with continuous dilution (*N* = 3000 cells, *T* = 300 hr, 10 replicates). The first-order perturbation result (grey solid line, Eq. 25) and Gaussian moment closure (solid coloured lines, Appendix D) are compared with simulation means (±1 SE). Model B (circles) remains at ⟨*n*⟩*/*⟨*r*⟩ = 1; Models D (squares) and E (diamonds) show enrichment increasing with *s*. The moment closure agrees with simulations within statistical error across the full range.

#### 3.2.11 Generalisation to nonlinear feedback functions

The preceding analysis assumed a linear dependence of translation rate on growth rate. Experimental evidence, however, shows that the relationship between ribosome abundance and growth rate is saturating rather than strictly linear (Dai et al., 2016). We now show that the main result generalises to an arbitrary functional form.

Let the translation rate per mRNA be given by a general smooth function *g*(*µ*) of the growth rate, so that the translation reaction is *m* → *m* + protein at rate *g*(*µ*(*n*)) · *m*. The linear case corresponds to *g*(*µ*) = *k*_*p*_+*q*_2_*µ*; a saturating (Michaelis–Menten) form would be *g*(*µ*) = *k*_*p*_+*q*_2_*µ/*(*K*_*µ*_+*µ*).

At zeroth order (*s* = 0), the growth rate is *µ*_0_ and the effective translation rate is 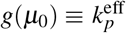, exactly as before. All zeroth-order moments, including the intrinsic covariance *C*_*nm*_, are unchanged.

At first order in *s*, we expand *g*(*µ*_0_+*sn*) ≈ *g*(*µ*_0_) + *g*^′^(*µ*_0_) *sn*, where *g*^′^(*µ*_0_) = *dg/dµ*|_*µ*0_. The perturbation calculation of Section 3.2.4 carries through with *g*^′^(*µ*_0_) replacing *q*_2_, yielding

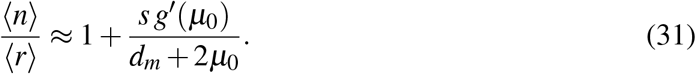

This is positive whenever *g*^′^(*µ*_0_) *>* 0, that is, whenever the translation rate is an increasing function of the growth rate at the operating point. The shape of the curve determines the magnitude of selective enrichment but not its existence.

For the saturating case *g*(*µ*) = *k*_*p*_+*q*_2_*µ/*(*K*_*µ*_+*µ*), the relevant derivative is

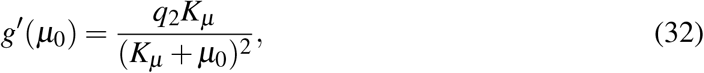

which is strictly positive for *q*_2_, *K*_*µ*_ *>* 0. The enrichment is weaker than in the linear case (since *g*^′^(*µ*_0_) *< q*_2_) but remains present. This is consistent with the computational finding of Ciechonska et al. (2022) that a saturating translation feedback produces selective enrichment (Ciechonska et al. 2022, Supplementary Material, Figure S10).

Theorem 2 (no enrichment under transcriptional feedback) also generalises to this setting, since the mRNA source ⟨*h*(*µ*(*n*))⟩ remains gene-independent for any transcription function *h*.

These generalisations extend beyond the perturbative level:

##### Theorem 3

(Exact covariance relation for selective enrichment). *Consider the two-stage model with an arbitrary fitness landscape µ*(*n*), *an arbitrary transcription function h*(*µ*), *and an arbitrary translation function g*(*µ*). *At steady state*,

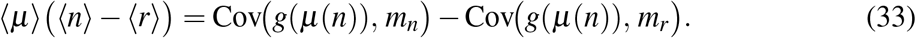

*This requires no perturbative expansion, no moment closure, and no linearity assumptions on µ*(*n*), *h*(*µ*), *or g*(*µ*). *Selective enrichment exists if and only if the translation rate covaries asymmetrically with the two mRNAs. In particular, Theorems 1 and 2 are recovered as corollaries: when g is constant (no translational feedback), both covariances vanish identically and* ⟨*n*⟩ = ⟨*r*⟩.

*Proof*. At steady state, the protein equations with the dilution–selection cancellation (Theorem 1) give ⟨*g*(*µ*(*n*)) *m*_*n*_⟩ = ⟨*µ*⟩⟨*n*⟩ and ⟨*g*(*µ*(*n*)) *m*_*r*_⟩ = ⟨*µ*⟩⟨*r*⟩. Since ⟨*m*_*n*_⟩ = ⟨*m*_*r*_⟩ for any transcription function *h*(*µ*) (Theorem 2), subtracting yields Eq. (33). The result reduces to Eq. (26) for linear *µ*(*n*) = *µ*_0_+*sn* and *g*(*µ*) = *k*_*p*_+*q*_2_*µ*.

Overall, the conditions for selective enrichment can be stated without reference to any particular functional form: selective enrichment occurs if and only if the translation rate is an increasing function of the growth rate, and the magnitude of enrichment is controlled by the local slope *g*^′^(*µ*_0_) of this relationship at the basal growth rate.

#### 3.2.12 Connection to the fluctuation–response relationship

The Fano factor of the fitness protein at *s* = 0 is

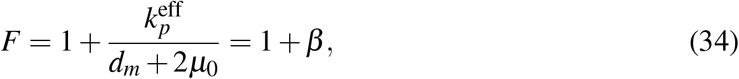

where 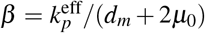 is the translational burst contribution. Let 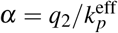 denote the growth-rate sensitivity of translation, normalised by the total effective translation rate. Then Eq. (25) can be rewritten as

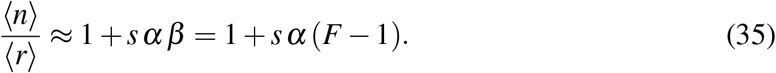

This has the structure of a fluctuation–response relation (Sato et al., 2003; Lehner and Kaneko, 2011): the response to selection (selective enrichment) is proportional to the fluctuation (*F* − 1) in the unperturbed system, modulated by the fraction *α* of translation that is growth-coupled. If gene expression is Poissonian (*F* = 1), there is no growth-driven enrichment.

This provides a direct analytical connection to the Ohm’s-law-like relationship between selection pressure and enrichment response observed by Ciechonska et al. (2022), where the “conductance” 1*/R* equals *αβ*. Figure 3 demonstrates this dependence across all model variants: enrichment increases monotonically with both the mRNA autocorrelation time *τ*_*m*_ = 1*/*(*d*_*m*_+*µ*_0_) (panel a) and the protein dilution time *τ*_*p*_ = 1*/µ*_0_ (panel b), while models without translational feedback remain at ⟨*n*⟩*/*⟨*r*⟩ = 1.

**Figure 3:**
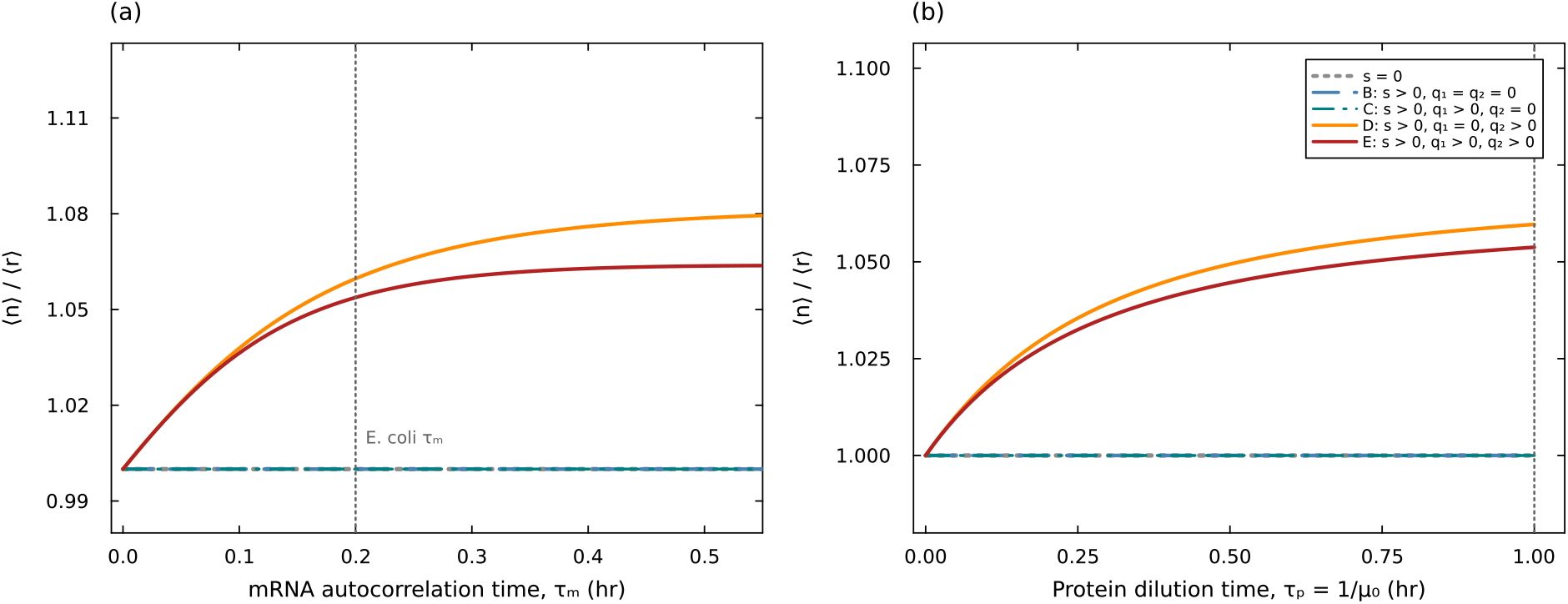
Enrichment ratio ⟨*n*⟩*/*⟨*r*⟩ as a function of two key timescales, at selection strength *s* = 0.15. **(a)** Ratio vs. mRNA autocorrelation time *τ*_*m*_ = 1*/*(*d*_*m*_+*µ*_0_) at fixed *µ*_0_ = 1. **(b)** Ratio vs. protein dilution time *τ*_*p*_ = 1*/µ*_0_ at fixed *d*_*m*_ = 4; as *µ*_0_ decreases (slower growth), both *τ*_*p*_ and *τ*_*m*_ increase, amplifying enrichment. In both panels, three model variants lacking translational feedback (no selection (*s* = 0), selection only (Model B), and selection with transcriptional feedback (Model C)) collapse to ⟨*n*⟩*/*⟨*r*⟩ = 1 (overlapping lines at unity). Only translational feedback (Models D and E) lifts the ratio above unity, with enrichment increasing monotonically with both *τ*_*m*_ and *τ*_*p*_. Model E (with transcriptional feedback, *q*_1_ *>* 0) is visibly below Model D (*q*_1_ = 0), showing that transcriptional feedback weakly suppresses selective enrichment by reducing intrinsic mRNA noise. Vertical dotted lines mark typical *E. coli* values (*τ*_*m*_ ≈ 0.2 hr, *τ*_*p*_ ≈ 1 hr). Parameters: *k*_*m*_ = 5, *k*_*p*_ = 3, *q*_1_ = 2, *q*_2_ = 3, computed from the Gaussian moment closure (Appendix D).

### 3.3 Concentration-dependent fitness

The preceding analysis assumed copy-number fitness: *µ*(*n*) = *µ*_0_+*sn*, where the growth rate depends on the total number of fitness protein molecules per cell. In many biological settings, however, the quantity that determines fitness is the protein concentration *n/V*, where *V* is the cell volume. In particular, the agent-based simulations of Ciechonska et al. (2022) compute fitness from the concentration of chloramphenicol acetyltransferase, and report concentrations as their output. We therefore extend the framework to concentration-dependent fitness:

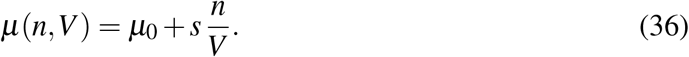

In a size-triggered division model, the cell volume grows deterministically within each cell cycle as *V* (*a*) = *e*^*µa*^ (normalising the birth volume to 1), with division occurring at *V* = 2, i.e. at the deterministic interdivision time *T* = ln 2*/µ*. In a steady-state population, whether exponentially growing or maintained at constant size (e.g., in a chemostat or Moran model), the age distribution is not uniform on [0, *T*] but exponential: younger cells are always more abundant because older cells have had more time to be randomly replaced. The steady-state age density is *p*(*a*) = 2*µ e*^−*µa*^ for *a* ∈ [0, *T*], obtained from the balance between ageing and removal with the boundary condition that each division produces two newborn cells. The population average of 1*/V* is therefore

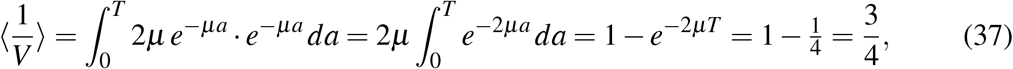

where we used *e*^−*µT*^ = 1*/*2. Interestingly, ⟨1*/V*⟩ is independent of the growth rate *µ*. This is because the exponential growth profile *V* = *e*^*µa*^ and the exponential age distribution combine to cancel the *µ*-dependence exactly.

With concentration-dependent fitness, the selection term in the general moment equation (3) becomes *s* Cov(*n/V, ϕ*) instead of *s* Cov(*n, ϕ*). At leading order in *s* (weak selection), the intracellular copy number *n* and cell volume *V* are approximately independent (*V* is deterministic at fixed growth rate, and stochastic variation in *n* perturbs *µ* only weakly). In this regime:

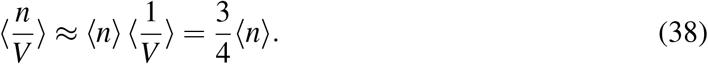

Consequently, the concentration fitness *µ* = *µ*_0_+*s*(*n/V*) is equivalent at leading order to copynumber fitness with a rescaled selection strength:

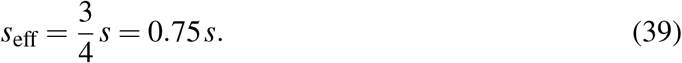

Theorems 1 and 2 are preserved under concentration-dependent fitness. For Model B, the cancellation between the dilution term ⟨*µ*(*n,V*) · *n*⟩ and the selection term *s* Cov(*n/V, n*) holds in the same algebraic form as in the copy-number case: both genes experience the same concentration-dependent growth rate, and the algebraic structure that ensures ⟨*n*⟩*/*⟨*r*⟩ = 1 is unchanged. The ratio remains exactly unity for any *s* ≥ 0. The same argument extends to Model C with transcriptional feedback only.

For Models D and E with translational feedback (*q*_2_ *>* 0), the first-order enrichment ratio under concentration-dependent fitness is obtained by replacing *s* with *s*_eff_ in Eq. (25):

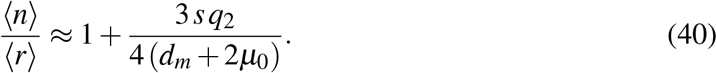

The geometric factor 3*/*4 reflects the cell-cycle averaging of volume variation: the concentration *n/V* is largest just after division (when *V* ≈ 1) and smallest just before (when *V* ≈ 2), and the average over the cycle reduces the effective selection by exactly 25% compared to the copynumber case. All structural conclusions carry through: selective enrichment requires *q*_2_ *>* 0, is independent of *q*_1_ at leading order, and scales with the super-Poisson noise (*F* − 1).

When selective enrichment is measured in concentrations rather than copy numbers, the relevant ratio is ⟨*n/V*⟩*/*⟨*r/V*⟩. At weak selection, *n* and *V* are approximately independent, so

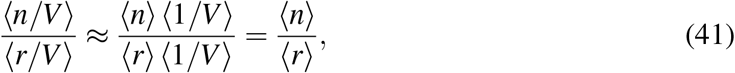

and the concentration-measured enrichment ratio equals the copy-number ratio. At stronger selection, corrections from Cov(*n*, 1*/V*) appear: cells with more fitness protein grow faster, have larger volumes on average, and thus lower concentrations, producing a negative covariance that slightly reduces the concentration ratio relative to the copy-number ratio. This effect enters at *O*(*s*^2^) and does not alter the leading-order result.

### 3.4 Growth rate recovery under selection

Selective enrichment not only shifts the relative abundance of fitness-gene transcripts but also increases the population mean growth rate. Since *µ*(*n*) = *µ*_0_+*sn*, the mean growth rate is ⟨*µ*⟩ = *µ*_0_+*s*⟨*n*⟩. Applying Eq. (3) to *ϕ* = *µ*(*n*) = *µ*_0_+*sn* gives

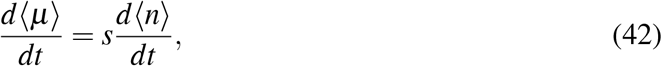

which, after expanding *d*⟨*n*⟩*/dt* using Eq. (3), contains a strictly positive selection contribution *s*^2^ Var(*n*) *>* 0. This is a direct manifestation of Fisher’s fundamental theorem (Fisher, 1930): the rate of increase in mean fitness is proportional to the variance in fitness, Var(*µ*) = *s*^2^ Var(*n*). For the linear fitness landscape *µ*(*n*) = *µ*_0_+*sn*, this means that phenotypic heterogeneity in the fitness gene (Var(*n*) *>* 0) drives the population mean growth rate upward.

#### 3.4.1 Application to histidine biosynthesis

In the histidine starvation experiments of Tsuru et al. (2011), the growth rate depends on the intracellular level of the biosynthetic enzyme HisC via a saturating response which can be modelled as

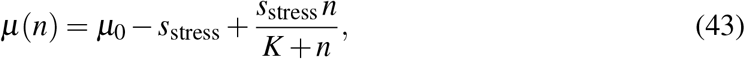

where *µ*_0_ is the unstressed growth rate, *s*_stress_ is the maximum growth rate penalty imposed by histidine depletion, and *K* is the half-saturation constant. Under full stress with no fitness protein (*n* = 0), the growth rate is *µ*_0_ − *s*_stress_; as *n* → ∞, the growth rate recovers to *µ*_0_.

In the linear regime (⟨*n*⟩ ≪ *K*), the approximation *n/*(*K*+*n*) ≈ *n/K* maps Eq. (43) onto the standard framework with a reduced basal growth rate *µ*_0_ − *s*_stress_ and effective selection strength *s*_eff_ = *s*_stress_*/K*. For Model B (growth-coupled dilution), the steady-state protein mean is ⟨*n*⟩ = *b/*(*µ*_0_ − *s*_stress_), giving the recovery condition

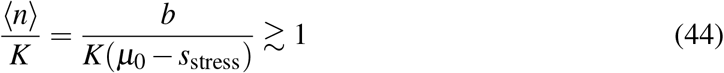

for substantial growth rate recovery. To put into words, the population growth rate recovers when the cell’s expression capacity (set by the ratio of the protein production rate *b* to the stressed dilution rate) exceeds the half-saturation constant *K* of the enzymatic pathway. With translational feedback (*q*_2_ *>* 0), selective enrichment further elevates ⟨*n*⟩ above the neutral level, broadening the parameter regime in which recovery occurs.

This result connects phenotypic selection to the emergence of drug resistance: even when phenotypic selection alone cannot fully restore the pre-stress growth rate, by partially recovering growth it extends the window during which cell division occurs under stress, increasing the opportunity for permanent genetic resistance mutations to arise.

### 3.5 Regulated expression bypasses the need for translational feedback

Theorems 1 and 2 assume constitutive gene expression. Many genes, however, are regulated by promoters that stochastically switch between active (ON) and inactive (OFF) states, often called the telegraph model of gene expression. We now show that such regulated expression provides an alternative route to selective enrichment under growth-coupled dilution, without requiring translational feedback (*q*_2_ *>* 0).

Consider a two-state promoter for each gene, with activation rate *k*_on_ and deactivation rate *k*_off_. Protein is produced at rate *b*_1_ when the promoter is ON and not at all when it is OFF; protein dilution is growth-coupled at rate *µ*(*n*) per molecule, as in Model B. Writing *g*_*n*_ ∈ {0, 1} for the fitness gene promoter state and *g*_*r*_ for the reference, the universal dilution–selection cancellation (Theorem 1) still holds, giving

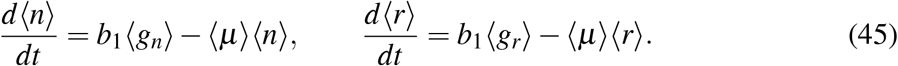

At steady state, the enrichment ratio reduces to

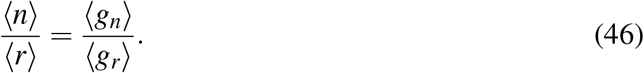

Enrichment thus arises entirely from a difference in the population-averaged promoter ON fractions. The moment equation for the fitness gene promoter state is

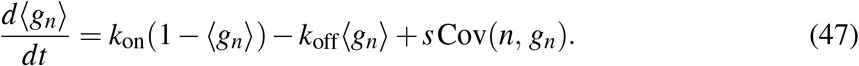

The selection term *s* Cov(*n, g*_*n*_) *>* 0 because, within a cell, having the fitness promoter ON leads to higher *n* and hence higher growth rate: the promoter state is informative about fitness. For the reference gene, the promoter state *g*_*r*_ is independent of *n* within a cell, so Cov(*n, g*_*r*_) = 0 and ⟨*g*_*r*_⟩ = *k*_on_*/*(*k*_on_+*k*_off_) (the neutral value).

At steady state, ⟨*g*_*n*_⟩ = (*k*_on_+*s* Cov(*n, g*_*n*_))*/*(*k*_on_+*k*_off_), giving

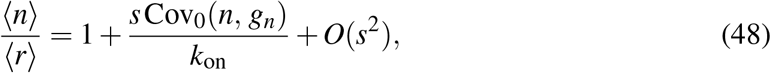

where Cov_0_ denotes the covariance evaluated at the neutral (*s* = 0) steady state. Computing Cov_0_(*n, g*_*n*_) from the standard telegraph model with dilution rate *µ*_0_ yields

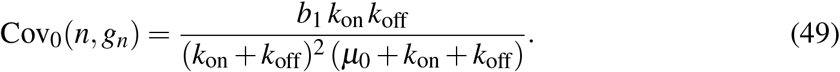

Let *F*_0_ = 1 + *b*_1_ *p*_off_*/*(*µ*_0_+*k*_*s*_) be the Fano factor of the neutral protein distribution, where *p*_off_ = *k*_off_*/*(*k*_on_+*k*_off_) and *k*_*s*_ = *k*_on_+*k*_off_ is the total switching rate. Substituting into Eq. (48):

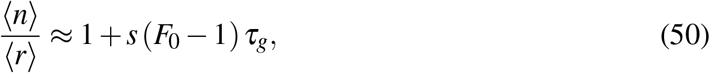

where *τ*_*g*_ = 1*/k*_*s*_ is the promoter autocorrelation time. This result has the same fluctuation– response structure as the translational feedback case (Eq. 35): enrichment is proportional to *s* times the super-Poissonian noise (*F*_0_ − 1), modulated by a memory timescale, here the promoter correlation time *τ*_*g*_ rather than the mRNA lifetime *τ*_*m*_. In the limit of fast switching (*k*_*s*_ → ∞), the Fano factor approaches unity, *τ*_*g*_ → 0, and Theorem 1 is recovered.

#### 3.5.1 Enrichment in the slow-switching regime

When the promoter switches slowly relative to protein turnover (*k*_*s*_ ≪ *µ*_0_), the protein distribution is bimodal: one mode near zero (promoter OFF) and one near *b*_1_*/µ*_0_ (promoter ON). In this regime, selection does not shift a single peak rightward, as in the translational feedback case, but instead changes the relative weights of the two modes. Cells with the fitness gene promoter ON have higher *n*, grow faster, and are overrepresented, increasing the weight of the upper mode for the fitness gene. The reference gene’s promoter, being uncorrelated with fitness, retains its neutral mode balance. The enrichment signature is therefore a shift in mode occupancy rather than a unimodal mean shift, consistent with the bimodal dynamics observed in agent-based simulations of regulated expression under selection (Ciechonska et al., 2022). Figure 4 confirms these predictions: at *s* = 0, both genes show identical bimodal distributions (panel a); under selection, the fitness gene’s upper mode gains weight while the reference gene retains its neutral mode balance (panel b); and the enrichment ratio matches both the first-order perturbation (Eq. 50) and an exact nonlinear solution (Appendix F) across selection strengths (panel c).

**Figure 4:**
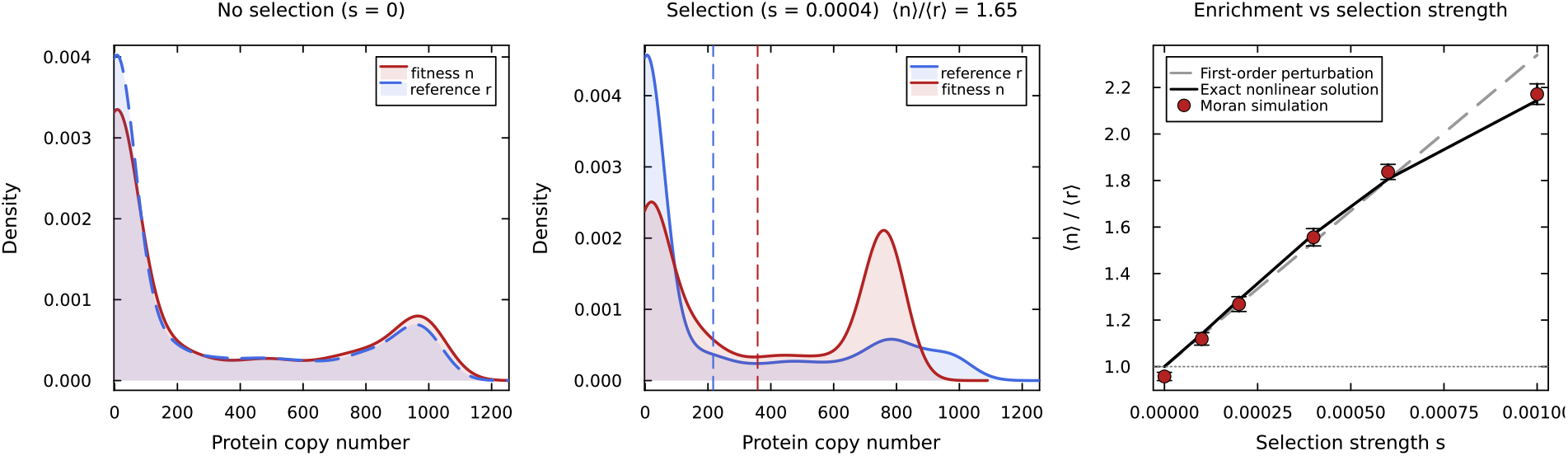
Selective enrichment from regulated expression under growth-coupled dilution, without translational feedback. **(a)** Neutral (*s* = 0): both genes show identical bimodal distributions from slow promoter switching (*k*_on_ = 0.1 hr^−1^, *k*_off_ = 0.3 hr^−1^, *b*_1_ = 1000 hr^−1^, *µ*_0_ = 1 hr^−1^). **(b)** Under selection (*s* = 4 × 10^−4^): the fitness gene’s upper mode gains weight while the reference gene retains its neutral mode balance. Dashed vertical lines mark the means. **(c)** Enrichment ratio vs. selection strength: first-order perturbation (Eq. 50, grey dashed), exact nonlinear solution (black solid; Appendix F), and Moran simulations (*N* = 3000 cells, *T* = 300 hr, 15 replicates ±1 SE). The Fano factor of the neutral distribution is *F*_0_ ≈ 536 and the promoter correlation time is *τ*_*g*_ = 2.5 hr.

The promoter-state selection mechanism (Cov(*n, g*_*n*_) *>* 0) operates equally under death-driven selection (Section 3.6), where it provides a quantitative enhancement to the enrichment that already arises from constitutive expression. The distinction is that for growth-driven selection, the dilution–selection cancellation eliminates the direct channel, making the promoter-state channel the sole route to enrichment; for death-driven selection, both channels are active.

### 3.6 Growth-driven versus death-driven selection

The preceding analysis establishes that growth-driven enrichment of constitutively expressed genes requires translational feedback (*q*_2_ *>* 0), or alternatively, a source of super-Poissonian gene-specific noise such as promoter switching (Section 3.5). However, experimental studies have reported selective enrichment without invoking any such mechanism. Lasri et al. (2020); Lasri and Sturrock (2021) observed phenotypic selection of MGMT-expressing glioblastoma cells under temozolomide (TMZ) stress, where MGMT confers resistance by repairing DNA damage and thereby reducing cell death. Similarly, the ampicillin model of Ciechonska et al. (2022) (Fig. 5 therein) includes a cell death rate that decreases with intracellular *β*-lactamase, rather than a growth rate that increases with the fitness protein. These are examples of death-driven selection, a framework that extends naturally to mammalian cells, where the bacterial coupling between growth rate and translational capacity (Klumpp et al., 2009; Dai et al., 2016) is not established, and to chemotherapy resistance, where drug-sensitive cells are selectively killed while cells that stochastically express higher levels of resistance-conferring proteins persist. Our moment framework reveals why such systems do not require translational feedback.

**Figure 5:**
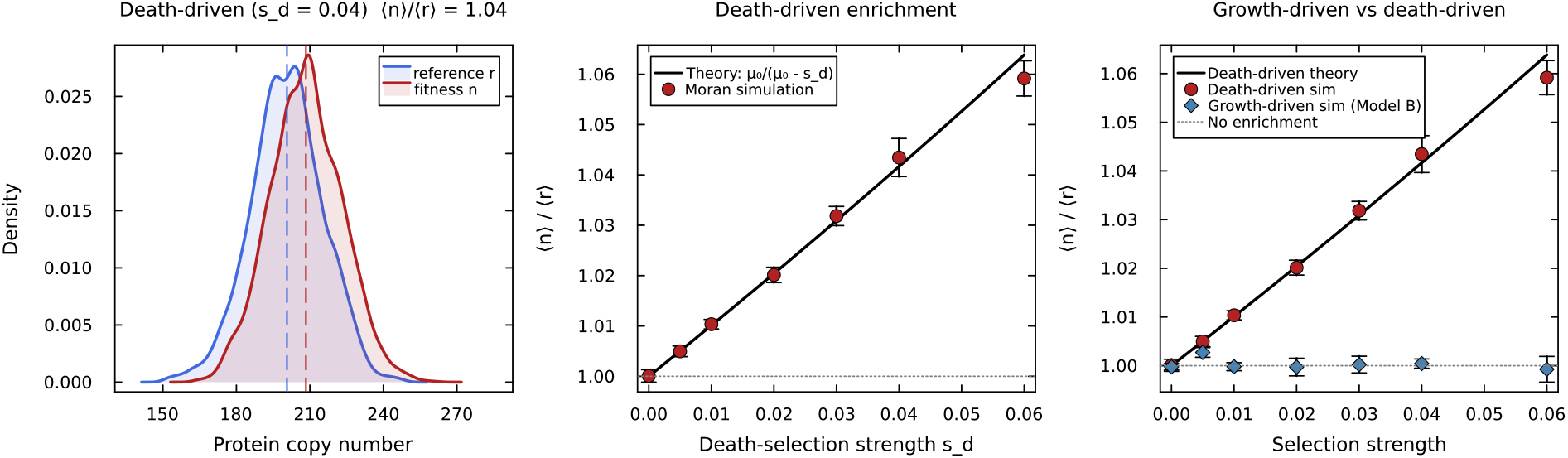
Death-driven versus growth-driven selective enrichment in a protein-only model without translational feedback. **(a)** Protein distributions under death-driven selection (*s*_*d*_ = 0.04, *γ*(*n*) = *γ*_0_ − *s*_*d*_*n*): the fitness gene (red) is shifted right relative to the reference (blue). **(b)** Enrichment ratio vs. death-selection strength: Moran simulations (*N* = 3000, *T* = 200 hr, 8 replicates ±1 SE) match the analytical prediction *µ*_0_*/*(*µ*_0_ − *s*_*d*_) (solid line). **(c)** Comparison: death-driven selection (red circles) produces enrichment that increases with selection strength, while growth-driven selection with growth-coupled dilution (blue diamonds, Model B) remains at ⟨*n*⟩*/*⟨*r*⟩ = 1 due to the exact dilution–selection cancellation. Parameters: *b* = 200 hr^−1^, *µ*_0_ = 1 hr^−1^, *γ*_0_ = 0.5 hr^−1^.

To see this, we generalise the total fitness to *w*(*n*) = *µ*(*n*) − *γ*(*n*), where *µ*(*n*) is the division rate and *γ*(*n*) is the death rate. Under the normalised master equation, the general moment equation for the mean fitness protein becomes

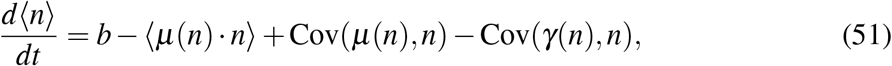

where *b* denotes the total protein production rate. The first selection-related term, −⟨*µ*(*n*) · *n*⟩, is the dilution penalty: faster-dividing cells dilute their proteins more rapidly. The second, Cov(*µ*(*n*), *n*), is the growth-driven selection benefit.

The dilution term and the growth selection term sum exactly:

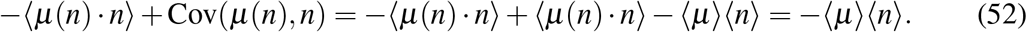

This is the same cancellation identified in Theorem 1: for purely growth-driven selection (*γ* = 0), the selective advantage of faster division is perfectly offset by the faster dilution of proteins. The mean equation closes to *d*⟨*n*⟩*/dt* = *b* − ⟨*µ*⟩⟨*n*⟩, and the fitness and reference genes experience identical dynamics, giving ⟨*n*⟩*/*⟨*r*⟩ = 1. This exact cancellation is the reason growth-driven enrichment requires translational feedback (*q*_2_ *>* 0): only a growth-rate-dependent translation rate, which multiplies the gene-specific mRNA, introduces an asymmetry between the two genes that survives the dilution–selection cancellation.

For death-driven selection, however, the situation differs. Consider the case where the growth rate is constant (*µ*(*n*) = *µ*_0_) and the death rate decreases with the fitness protein: *γ*(*n*) = *γ*_0_ *h*(*n*) with *h*^′^(*n*) *<* 0. This describes the Lasri TMZ model, where *γ*(*n*) ∝ [TMZ]*/*([TMZ] + *s* · *n*), and the Ciechonska ampicillin model, where intracellular ampicillin (which induces death) is cleaved by the fitness protein. The moment equation becomes

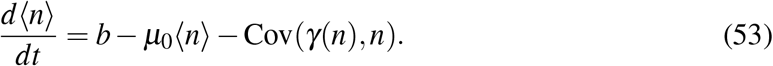

The dilution term is now simply −*µ*_0_⟨*n*⟩ (constant, gene-independent), and the growth-selection term vanishes because *µ* is constant. The only selection contribution is −Cov(*γ*(*n*), *n*).

Because the death rate *γ*(*n*) decreases with *n* (cells with more fitness protein are less likely to die), the covariance Cov(*γ*(*n*), *n*) is strictly negative. Therefore −Cov(*γ*(*n*), *n*) *>* 0: death-driven selection acts as a positive source term that raises ⟨*n*⟩ above its unselected value. This source term carries no corresponding dilution penalty. Escaping death does not force a cell to dilute its proteins faster; it simply allows the cell and its protein content to persist. The mean equation for the reference protein is *d*⟨*r*⟩*/dt* = *b* − *µ*_0_⟨*r*⟩ (no covariance term, since *γ* depends on *n*, not on *r*), giving ⟨*r*⟩ = *b/µ*_0_ at steady state. The fitness protein, by contrast, satisfies ⟨*n*⟩ = *b/µ*_0_ −Cov(*γ*(*n*), *n*)*/µ*_0_ *>* ⟨*r*⟩, and the enrichment ratio exceeds unity.

This mechanism is purely noise-driven: baseline stochastic gene expression generates a distribution of protein levels, and cells in the low-*n* tail of this distribution die preferentially, shifting the population mean upward. No translational feedback, no explicit mRNA dynamics, and no growth-rate coupling are required. For a linear death rate *γ*(*n*) = *γ*_0_ − *s*_*d*_*n* with Poisson protein statistics, the enrichment ratio is ⟨*n*⟩*/*⟨*r*⟩ = *µ*_0_*/*(*µ*_0_ − *s*_*d*_), analogous to the Mora–Walczak result (Eq. 5) but with the constant dilution rate *µ*_0_ replacing the active degradation rate *d*. This explains why the protein-only models of Lasri et al. (2020); Lasri and Sturrock (2021) and the ampicillin model of Ciechonska et al. (2022) produce selective enrichment without requiring translational feedback, explicit mRNA dynamics, or growth-rate coupling: death-driven selection arises naturally from the overlap between gene expression noise and a survival threshold. Figure 5 confirms these predictions: Moran simulations show that death-driven selection produces enrichment (panel a,b) matching the analytical prediction, while growth-driven selection with growth-coupled dilution (Model B) produces no enrichment at any selection strength (panel c).

When explicit mRNA dynamics and growth-related feedbacks are included in a death-driven model, the enrichment magnitude changes but its existence does not: translational feedback (*q*_2_ *>* 0) enhances selective enrichment by increasing the translational burst size 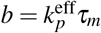 and hence the protein noise, while transcriptional feedback (*q*_1_) remains absent at first order in the selection strength, as in the growth-driven case.

Therefore, the requirement for translational feedback (*q*_2_ *>* 0) is specific to growth-driven enrichment of constitutively expressed genes, where the exact dilution–selection cancellation must be broken by a gene-specific mechanism. Death-driven enrichment operates through a distinct pathway (selective removal of low-expressing cells) that bypasses this cancellation entirely. Regulated expression via promoter switching (Section 3.5) enhances death-driven enrichment by providing an additional promoter-state channel (Cov(*n, g*_*n*_) *>* 0), but is not required for it: the direct survival advantage of high-expressing cells is sufficient. The general moment framework (Eq. 51) unifies all three mechanisms (translational feedback, promoter-state selection, and death-driven selection) and clarifies when each set of minimal ingredients applies.

## 4 Discussion

We have provided an analytical framework for phenotypic selection of fitness-conferring genes, identifying the conditions under which selective enrichment arises across growth-driven and death-driven selection regimes, for both constitutive and regulated gene expression. Computational screening (Ciechonska et al., 2022) identified the minimal ingredients for selective enrichment by testing ten model variants; the analytical framework developed here reveals the mechanistic basis for each requirement, explains alternative routes to enrichment (regulated expression, death-driven selection) within the same framework, and yields quantitative predictions that are difficult to extract from simulation alone.

For constitutively expressed genes under growth-driven selection, the no-enrichment theorems (Theorems 1 and 2), the exact covariance relation (Theorem 3), and the enrichment formula (25) together constitute a design rule: selective enrichment requires a growth-rate-dependent feedback that acts on a rate which multiplies a gene-specific stochastic variable, and the resulting response scales with the super-Poisson noise of the unperturbed circuit (Eq. 35). We now discuss the mechanistic basis of these results.

The failure of transcriptional feedback to produce selective enrichment can be understood in terms of timescales and memory. In the two-stage model, each mRNA molecule lives for a characteristic time *τ*_*m*_ = 1*/*(*d*_*m*_+*µ*_0_) before being degraded or diluted. During its lifetime, each mRNA produces a burst of proteins, and the autocorrelation time of mRNA fluctuations sets a memory window over which a cell retains information about which gene happened to be transcribed more actively. Transcriptional feedback enhances the rate of mRNA production, but this rate is gene-independent: the feedback-enhanced transcription rate *k*_*m*_+*q*_1_*µ*(*n*) is identical for the fitness gene and the reference gene within the same cell, so that both genes receive the same boost when the growth rate increases. This ensures ⟨*m*_*n*_⟩ = ⟨*m*_*r*_⟩ exactly (Eq. 12), and the downstream protein dynamics inherit this symmetry.

Translational feedback, by contrast, acts at a stage where gene-specific stochastic memory exists. The key timescale is the mRNA autocorrelation time

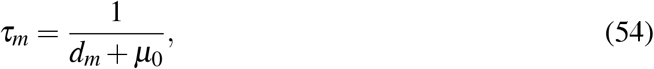

which sets the duration over which a protein remains correlated with the mRNA molecule that produced it. In the two-stage model, the intrinsic protein–mRNA covariance is

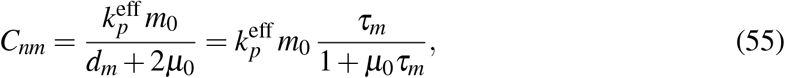

which is positive and increases with *τ*_*m*_: longer-lived mRNAs create stronger correlations between a protein and its own mRNA. The fitness protein *n* is correlated with its own mRNA *m*_*n*_ but uncorrelated with the reference mRNA *m*_*r*_ at *s* = 0. The translation rate (*k*_*p*_+*q*_2_*µ*(*n*)) · *m* multiplies the gene-specific mRNA copy number *m*, so a cell that has, by chance, more *m*_*n*_ than *m*_*r*_ will translate proportionally more protein *n* when the growth rate is elevated. The resulting excess of protein *n* persists on the much longer protein timescale *τ*_*p*_ = 1*/µ*_0_ ≫ *τ*_*m*_, maintaining the selective advantage across multiple mRNA lifetimes and cell division events. This produces a positive feedback loop (more *n* leads to faster growth, which increases translation of *m*_*n*_, which produces more *n*) that does not exist for the reference gene.

The enrichment ratio (Eq. 25) can be rewritten in terms of these timescales as

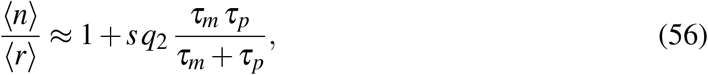

since 1*/*(*d*_*m*_ + 2*µ*_0_) = 1*/*(1*/τ*_*m*_ + 1*/τ*_*p*_) = *τ*_*m*_*τ*_*p*_*/*(*τ*_*m*_+*τ*_*p*_). The memory effect is thus simply proportional to the harmonic mean of the mRNA and protein timescales, which makes explicit that selective enrichment requires both finite mRNA memory (*τ*_*m*_ *>* 0, i.e. explicit mRNA dynamics) and translational feedback (*q*_2_ *>* 0) to exploit that memory. In the limit *τ*_*m*_ → 0 (instantaneous mRNA, equivalent to a protein-only model), *C*_*nm*_ → 0 and enrichment vanishes, consistent with the failure of Models A and B. In the limit *τ*_*p*_ → 0 (infinitely fast dilution), enrichment also vanishes because protein fluctuations are immediately washed out. This dependence on *τ*_*m*_ is shown in Figure 3(a); the complementary dependence on *τ*_*p*_ = 1*/µ*_0_ (the protein dilution timescale) is shown in Figure 3(b). Active protein degradation (rate *d*_*p*_) would reduce the effective protein timescale to *τ*_*p*_ = 1*/*(*µ*_0_+*d*_*p*_), further suppressing enrichment. Most bacterial proteins are highly stable and are diluted almost entirely by growth rather than active proteolysis, which maximises *τ*_*p*_ and hence enrichment. Proteins subject to active degradation, such as SsrA-tagged substrates of the ClpXP protease, would be predicted to show weaker selective enrichment, all else being equal.

These timescale arguments lead to a general principle for constitutive growth-driven enrichment: selective enrichment requires a growth-rate-dependent feedback that acts on a rate which multiplies a gene-specific stochastic variable, creating asymmetric noise amplification on a timescale long enough to persist across dilution events. Transcription rate enhancement does not satisfy this condition, since it does not multiply a gene-specific stochastic variable. Translation rate enhancement does, since it multiplies the gene-specific mRNA *m*, and the resulting protein fluctuations persist on the slow timescale *τ*_*p*_.

The regulated expression analysis (Section 3.5) reveals a broader principle: any source of gene-specific noise with sufficient memory can drive enrichment, even without translational feedback. In the two-state promoter model, the promoter state *g*_*n*_ is a gene-specific stochastic variable with correlation time *τ*_*g*_ = 1*/*(*k*_on_+*k*_off_). Selection enriches for cells where the fitness gene promoter is ON (Cov(*n, g*_*n*_) *>* 0) but cannot distinguish the reference promoter state (Cov(*n, g*_*r*_) = 0), producing enrichment ∝ *s*(*F*_0_ − 1)*τ*_*g*_. This unifies the two mechanisms: translational feedback exploits mRNA memory (*τ*_*m*_), while regulated expression exploits promoter-state memory (*τ*_*g*_). Both produce enrichment proportional to the super-Poissonian noise (*F* − 1) times the relevant correlation time. Transcriptional bursting (Golding et al., 2005; Raj et al., 2006) amplifies enrichment in both cases: under translational feedback, bursts inflate the mRNA variance and hence *C*_*nm*_; under regulated expression, slow promoter switching directly increases *F*_0_ and *τ*_*g*_. The regulated expression mechanism is closely related to bet-hedging (Balaban et al., 2004; Kussell and Leibler, 2005): stochastic promoter switching generates pre-existing phenotypic heterogeneity that selection exploits by differentially favouring one promoter state.

The linear relationship between selection pressure and enrichment response observed by Ciechonska et al. (2022) *is a direct consequence of Eq. (25): at weak selection, the ratio is linear in s* with slope *q*_2_*/*(*d*_*m*_ + 2*µ*_0_). The conductance in the Ohm’s law analogy of Ciechonska et al. (2022) is precisely this slope, determined by the noise properties of the gene expression circuit. The connection to the fluctuation–response relationship (Sato et al., 2003; Lehner and Kaneko, 2011) is made explicit by Eq. (35): selective enrichment is proportional to the super-Poisson component of protein noise, *F* − 1, where *F* is the Fano factor of the unperturbed (*s* = 0) gene expression distribution. This is an experimentally actionable prediction: one could measure the Fano factor of a gene under non-selective conditions and, given an estimate of the growthrate coupling 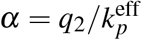, predict the magnitude of selective enrichment under stress without free parameters. If gene expression were Poissonian (*F* = 1), there would be no growth-driven enrichment regardless of how strong the selection or the translational feedback.

More broadly, the exact relation (26) provides a level of mechanistic insight that simulations and model selection alone cannot. Rewriting it as

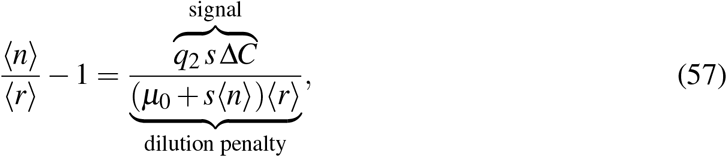

decomposes selective enrichment into a signal (translational feedback times selection strength times gene-specific covariance asymmetry) divided by a dilution penalty that grows with protein abundance. Growth-coupled dilution is not itself an ingredient for enrichment; rather, it is the biologically realistic feature that creates the exact cancellation (Theorem 1), making enrichment impossible without an additional mechanism to break the symmetry. The signal-to-penalty decomposition explains what that mechanism must provide: explicit mRNA dynamics are required because Δ*C* is built from the intrinsic protein–mRNA covariance *C*_*nm*_, which vanishes in the protein-only limit (*τ*_*m*_ → 0), and translational feedback (*q*_2_ *>* 0) is required because it is the prefactor of the entire signal.

A positive coupling between growth rate and transcription rate is well established in bacterial physiology, arising from the growth-rate dependence of free RNA polymerase concentration (Klumpp and Hwa, 2008; Klumpp et al., 2009). The signal-to-penalty decomposition shows that this feedback not only cannot produce selective enrichment on its own (Theorem 2), but actively hinders it: transcriptional feedback raises the mean protein level (*n*_0_ scales with *k*_*m*_+*q*_1_*µ*_0_), inflating the dilution penalty in the denominator without proportionally strengthening the gene-specific signal in the numerator. The second-order expansion (Eq. 30) makes this precise: the leading *q*_1_-dependent correction is −*s*^2^*q*_2_*n*_0_*/*[*µ*_0_(*d*_*m*_ + 2*µ*_0_)], which is strictly negative and grows with *q*_1_ through *n*_0_. Both the exact relation and the Gaussian moment closure confirm that this increased dilution cost dominates any effect *q*_1_ may have on the covariance asymmetry Δ*C*; accordingly, Model E (*q*_1_ *>* 0, *q*_2_ *>* 0) produces weaker enrichment than Model D (*q*_1_ = 0, *q*_2_ *>* 0) at all selection strengths, as confirmed by Moran simulations (Figure 2). Importantly, however, the opposition is quantitative, not qualitative: the translational signal is strong enough to overcome the dilution penalty, and selective enrichment persists in the presence of biologically realistic transcriptional feedback. This is consistent with the Bayesian model selection of Ciechonska et al. (2022), which identified Model 7 (translational feedback only) as most likely to produce selective enrichment, with Model 8 (transcriptional and translational feedback) and Model 10 (regulated expression) also producing enrichment. Our analysis shows that the leading-order enrichment ratio is independent of transcriptional feedback strength, and that transcriptional feedback slightly suppresses enrichment at higher orders. The Ciechonska models use a saturating (Michaelis–Menten) fitness function, which can be linearised around the self-consistent operating point to give an effective selection strength *s*_eff_ that substitutes for *s* in Eq. (25), preserving all structural results (see Appendix D for details).

The extension to concentration-dependent fitness *µ* = *µ*_0_+*s*(*n/V*) (Section 3.3) preserves all structural results but reduces the enrichment magnitude by a parameter-free geometric factor of 3*/*4. This factor arises from averaging 1*/V* over the cell cycle: exponential volume growth combined with the exponential age distribution of a steady-state population yields ⟨1*/V*⟩ = 3*/*4, independent of the growth rate. The result is a 25% reduction in enrichment for any gene whose fitness depends on concentration rather than copy number, and connects directly to the agent-based model of Ciechonska et al. (2022), where fitness depends on the concentration of chloramphenicol acetyltransferase. Again, this is a structural prediction: it depends on no model parameters and would be difficult to identify from numerical simulations alone, which would show a quantitative reduction but not directly reveal its origin or universality.

Our analysis uses a continuous dilution approximation rather than explicit cell division with binomial partitioning. In Appendix E, we show that this approximation is exact at the mean level: Theorems 1–3 and all structural conclusions hold under discrete binomial partitioning. Figure 2 confirms these predictions against Moran simulations across the full range of selection strengths, using both the first-order perturbation result and the non-perturbative Gaussian moment closure (Appendix D). Theorems 1 and 2 hold for all *s*, and the exact relation (26) for Model E likewise requires no weak-selection assumption. For stronger selection, higher-order terms in *s* contribute appreciably to the enrichment ratio, but the structural result (selective enrichment of constitutively expressed genes requires *q*_2_ *>* 0) is unchanged.

The growth rate recovery analysis (Section 3.4) connects selective enrichment to a clinically relevant outcome: the population mean growth rate increases at a rate *s*^2^Var(*n*) (Fisher’s fundamental theorem), and for saturating fitness functions the degree of recovery is governed by the ratio ⟨*n*⟩*/K* of protein expression to the enzymatic half-saturation constant. Even partial recovery extends the window for cell division under stress, providing opportunities for permanent genetic resistance mutations to arise. This positions phenotypic selection as a potential early step in the emergence of drug resistance, in both bacterial antibiotic treatment and chemotherapy in mammalian cells.

## Acknowledgements

We thank Vahid Shahrezaei and Mark Isalan for discussions on the computational modelling that motivated this work.

## A Derivation of the general moment equation

Starting from Eq. (2), multiply both sides by *ϕ* (**x**) and sum over all states:

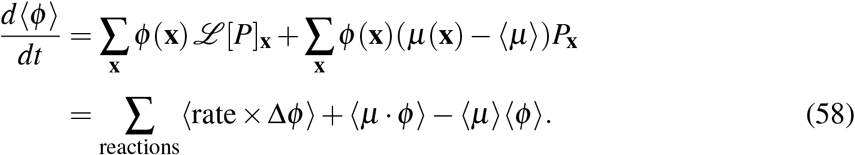

For linear selection *µ*(*n*) = *µ*_0_+*sn*, the selection term becomes *s*(⟨*nϕ*⟩ −⟨*n*⟩⟨*ϕ*⟩) = *s* Cov(*n, ϕ*).

## B Derivation of the protein–mRNA covariance

At *s* = 0, the two-stage model for a single gene has effective rates 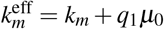 and 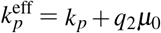. The steady-state means are 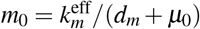 and 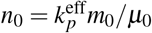. The Jacobian of the drift vector 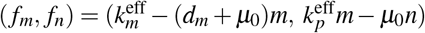 evaluated at (*m*_0_, *n*_0_) is

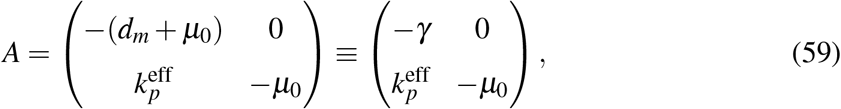

where *γ* = *d*_*m*_+*µ*_0_. The diffusion matrix (sum of production and turnover contributions) is

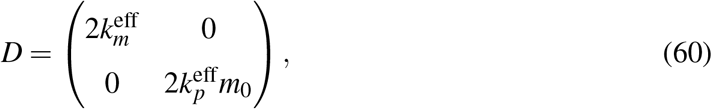

where the diagonal entries are 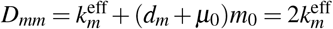 (using 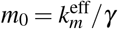) and 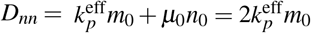 (using 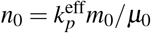).

The covariance matrix ∑ satisfies the Lyapunov equation *A*∑ + ∑*A*^*T*^+*D* = 0. Writing this out component by component:

The (*m, m*) entry gives *A*_11_∑_*mm*_ + ∑_*mm*_*A*_11_+*D*_*mm*_ = 0, i.e.

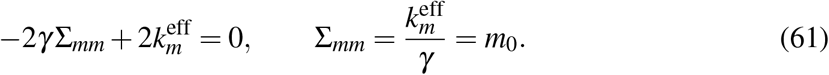

The (*m, n*) entry gives *A*_11_∑_*mn*_+*A*_21_∑_*mm*_ + ∑_*mn*_*A*_22_+*D*_*mn*_ = 0, i.e.

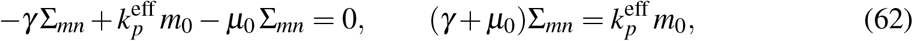

giving

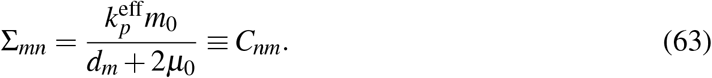

The (*n, n*) entry gives *A*_21_∑_*mn*_+*A*_22_∑_*nn*_ + ∑_*nn*_*A*_22_ + ∑_*mn*_*A*_21_+*D*_*nn*_ = 0, i.e.

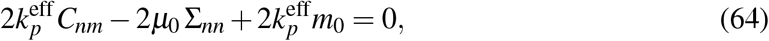

giving

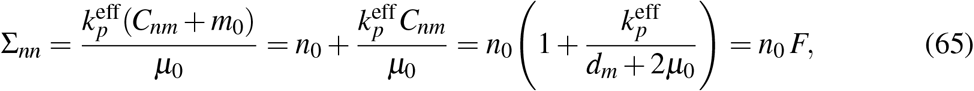

where 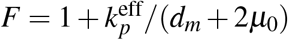 is the Fano factor.

## C Detailed perturbative calculation

We expand all quantities in powers of *s*:

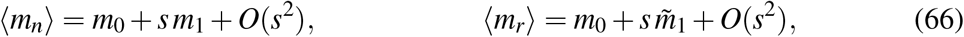

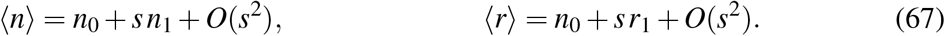

At *s* = 0, the fitness protein *n* and its own mRNA *m*_*n*_ are correlated, but *n* and the reference mRNA *m*_*r*_ are independent:

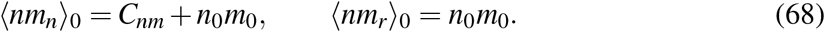

### C.0.1 mRNA equation at *O*(*s*)

The steady-state mRNA equation is 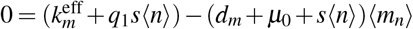. Substituting the expansions and collecting the *O*(*s*) terms yields:

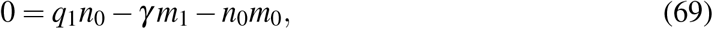

where *γ* = *d*_*m*_+*µ*_0_. Here the *q*_1_*n*_0_ comes from the transcriptional feedback (*q*_1_*s*⟨*n*⟩ at zeroth order in ⟨*n*⟩), the −*γ m*_1_ from total mRNA turnover (degradation plus dilution) acting on the first-order mRNA correction, and −*n*_0_*m*_0_ from the growth-enhanced mRNA decay (−*s*⟨*n*⟩⟨*m*_*n*_⟩ at zeroth order). Solving:

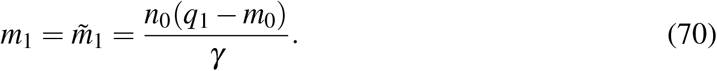

The first-order correction *m*_1_ is identical for both genes because the transcription rate depends on the growth rate (via *n*), which is gene-independent. Consequently *q*_1_ cannot produce genespecific effects.

#### C.0.2 Fitness protein equation at *O*(*s*)

The steady-state protein equation (Eq. 13) at all orders is

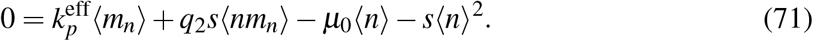

The *O*(1) balance gives 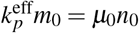, confirming 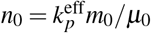. Writing this out component by component at *O*(*s*) yields:

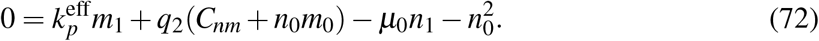

Note that *q*_2_ multiplies the zeroth-order cross-moment ⟨*nm*_*n*_⟩_0_ directly; there is no 1*/µ*_0_ prefactor.

Using 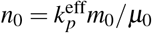, the *n*_0_*m*_0_ terms simplify:

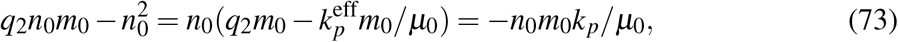

where we used 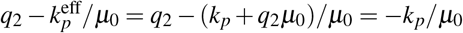. Thus:

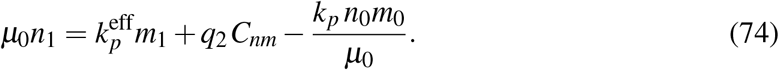

#### C.0.3 Reference protein equation at *O*(*s*)

From Eq. (14), using ⟨*nm*_*r*_⟩_0_ = *n*_0_*m*_0_ (since *n* and *m*_*r*_ are independent at *s* = 0):

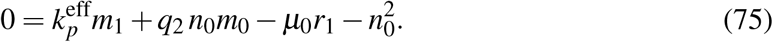

The only difference from Eq. (72) is the replacement of *C*_*nm*_+*n*_0_*m*_0_ by *n*_0_*m*_0_: the intrinsic covariance *C*_*nm*_ is absent because *n* and *m*_*r*_ are uncorrelated at *s* = 0. Using the same simplification:

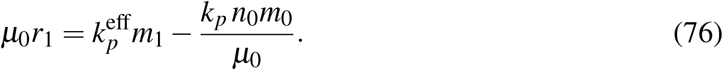

#### C.0.4 Enrichment ratio

Subtracting Eq. (75) from Eq. (72) yields

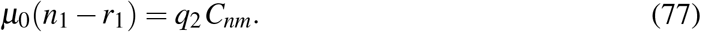

The common terms (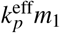 and −*k*_*p*_*n*_0_*m*_0_*/µ*_0_) cancel exactly, leaving only the intrinsic covariance term. Solving for the difference yields:

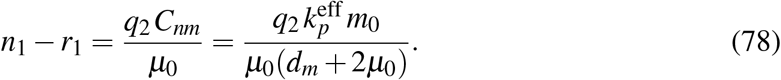

Dividing by 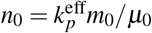:

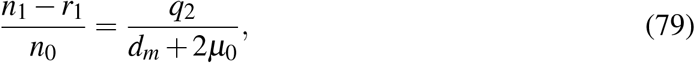

yielding the main result Eq. (25). Note that *q*_1_ does not appear: it enters both *n*_1_ and *r*_1_ identically through *m*_1_, and cancels in the difference.

## D Gaussian moment closure

The perturbation expansion of Appendix C is valid when the dimensionless parameter *sn*_0_*/µ*_0_ is small. To extend the analytical predictions to stronger selection, we truncate the moment hierarchy at second order by setting all third cumulants to zero (Gaussian closure). This yields 14 coupled algebraic equations at steady state: 4 for the means (⟨*m*_*n*_⟩, ⟨*n*⟩, ⟨*m*_*r*_⟩, ⟨*r*⟩) and 10 for the covariances *σ*_*i j*_ = Cov(*x*_*i*_, *x* _*j*_).

The mean equations follow from Eqs. (11)–(14), after applying the ⟨*n*^2^⟩–Var(*n*) cancellation of Theorem 1. Only two covariances enter the mean equations: Cov(*n, m*_*n*_) in the equation for ⟨*n*⟩ and Cov(*n, m*_*r*_) in the equation for ⟨*r*⟩, both multiplied by *q*_2_*s*. The covariance equations are obtained from the eight intracellular reactions using *dσ*_*i j*_*/dt* = *d*⟨*x*_*i*_*x* _*j*_⟩*/dt* − *µ*_*i*_ *dµ*_*j*_*/dt* − *µ*_*j*_ *dµ*_*i*_*/dt*, where each reaction contributes ⟨*w* · Δ(*x*_*i*_*x* _*j*_)⟩ − *µ*_*i*_⟨*w* · *νδ*_*jp*_⟩ − *µ*_*j*_⟨*w* · *νδ*_*ip*_⟩. Under the Gaussian closure, the selection term vanishes from the covariance equations. To see why, note that the selection contribution to *d*Cov(*X,Y*)*/dt* is

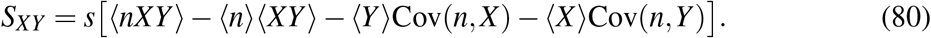

Expanding ⟨*nXY*⟩ in terms of central moments shows that *S*_*XY*_ = *sκ*_3_(*n, X,Y*), where *κ*_3_ is the third cumulant. Setting all third cumulants to zero (the definition of Gaussian closure) therefore eliminates the selection contribution identically, so that selection does not appear explicitly in the covariance equations.

We solve the 14 equations by Newton’s method with parameter continuation from the exact *s* = 0 solution (obtained from the Lyapunov equation of Appendix B), using an adaptive step size in *s* and a numerical Jacobian. The closure preserves three structural properties at every value of *s*: (i) ⟨*m*_*n*_⟩ = ⟨*m*_*r*_⟩ (mRNA equality); (ii) the exact relation (26); and (iii) ⟨*n*⟩*/*⟨*r*⟩ = 1 when *q*_2_ = 0. Property (iii) follows because the covariance equations decouple from the mean equations when *q*_2_ = 0, so the mean equations reduce to those of Model B. At weak selection the closure reproduces the first-order perturbation expansion; at stronger selection it provides a non-perturbative prediction that agrees with Moran simulations (Figure 2).

A notable prediction of the closure is that Model E (*q*_1_ *>* 0, *q*_2_ *>* 0) produces slightly weaker enrichment than Model D (*q*_1_ = 0, *q*_2_ *>* 0). This is explained analytically by the second-order correction (Eq. 28): the nonlinear dilution factor 1*/*(1 + *sn*_0_*/µ*_0_) suppresses enrichment when the protein levels are large, and since transcriptional feedback raises the zeroth-order protein level (*n*_0_ ∝ *k*_*m*_+*q*_1_*µ*_0_), it amplifies this dilution penalty without proportionally strengthening the gene-specific signal in the numerator. This suppression is confirmed by the Moran simulations at all selection strengths tested.

## E Equivalence of continuous and discrete dilution at the mean level

Consider a Moran model in which cell *i* divides at rate *µ*(*n*_*i*_) = *µ*_0_+*sn*_*i*_, molecules are partitioned binomially (each molecule independently assigned to one daughter with probability 1*/*2), and a randomly chosen cell is killed to maintain constant population size. In this discrete-dilution framework, the contribution of division to the population mean of any intracellular quantity *ϕ* decomposes into two terms: (i) the self-dilution of the dividing cell, −(1*/*2)⟨*µ*(*n*) ·*ϕ*⟩, where the factor 1*/*2 reflects the expected binomial loss; and (ii) the biased replacement, +(1*/*2)⟨*µ*(*n*) · *ϕ*⟩ −⟨*µ*⟩⟨*ϕ*⟩, where the second daughter replaces a random cell. These combine to give

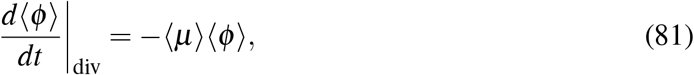

which is identical to the continuous-dilution result. The self-dilution loss and the biased-replacement gain cancel exactly, leaving only dilution at the population-mean growth rate, −⟨*µ*⟩⟨*ϕ*⟩, plus the selection contribution that is already captured by the ⟨*n*^2^⟩–Var(*n*) cancellation.

Consequently, Theorems 1–3 hold without modification in the discrete-dilution framework, since they follow entirely from the mean equations. In particular, the structural conclusion that selective enrichment requires *q*_2_ *>* 0 and is independent of *q*_1_ at leading order is unchanged. The continuous-dilution approximation does, however, affect the variance and covariance equations: binomial partitioning adds extra noise at each division event and alters the effective decay rate of correlations, modifying the single-cell steady-state covariance *C*_*nm*_. Since the firstorder enrichment ratio depends on *C*_*nm*_ (through 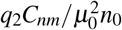), the specific numerical coefficient 1*/*(*d*_*m*_ + 2*µ*_0_) in Eq. (25) is particular to the continuous-dilution model; the Moran model with binomial partitioning would yield a quantitatively different coefficient, though the structural dependence on *q*_2_, *d*_*m*_, and *µ*_0_ is preserved. The Moran simulations used for validation in Figure 2 employ continuous intracellular dilution, and confirm the analytical formulas in that setting.

## F Exact nonlinear solution for regulated expression

We derive the nonlinear (all-orders-in-*s*) enrichment ratio for the two-state promoter model with growth-coupled dilution (Section 3.5). The intracellular state of each cell includes the promoter states *g*_*n*_, *g*_*r*_ ∈ {0, 1} (for the fitness and reference genes) and the protein copy numbers *n, r*. Protein is produced at rate *b*_1_*g*_*n*_ (or *b*_1_*g*_*r*_) and diluted at rate *µ*(*n*) = *µ*_0_+*sn* per molecule. The promoter activates at rate *k*_on_ and deactivates at rate *k*_off_, with *k*_*s*_ = *k*_on_+*k*_off_.

### Mean equations

The universal dilution–selection cancellation (Theorem 1) applies to any production process, giving the mean equations

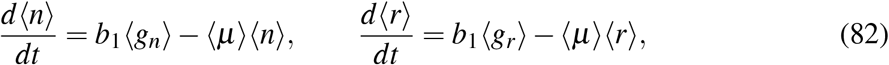

where ⟨*µ*⟩ = *µ*_0_+*s*⟨*n*⟩. At steady state, ⟨*n*⟩*/*⟨*r*⟩ = ⟨*g*_*n*_⟩*/*⟨*g*_*r*_⟩.

The promoter dynamics for the fitness gene give

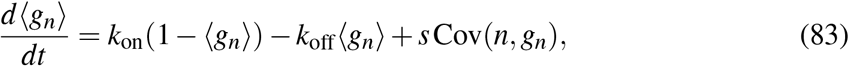

while for the reference gene, Cov(*n, g*_*r*_) = 0 (the reference promoter is independent of the fitness protein within each cell), so ⟨*g*_*r*_⟩ = *k*_on_*/k*_*s*_ at steady state.

### Cross-moment equation

To compute Cov(*n, g*_*n*_) = ⟨*ng*_*n*_⟩ − ⟨*n*⟩⟨*g*_*n*_⟩, we derive the moment equation for ⟨*ng*_*n*_⟩. Setting *ϕ* = *ng*_*n*_ in the general moment equation, the reaction contributions are:

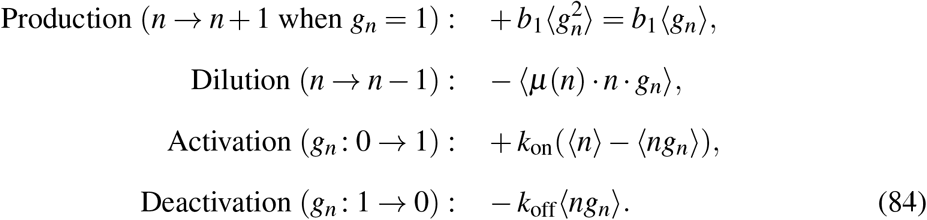

The selection contribution is *s* Cov(*n, ng*_*n*_) = *s*(⟨*n*^2^*g*_*n*_⟩ − ⟨*n*⟩⟨*ng*_*n*_⟩). The dilution term expands as −*µ*_0_⟨*ng*_*n*_⟩ − *s*⟨*n*^2^*g*_*n*_⟩. The ⟨*n*^2^*g*_*n*_⟩ terms from dilution and selection cancel exactly (another instance of the universal cancellation), yielding

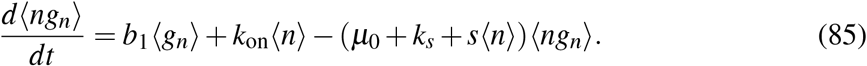

At steady state, writing 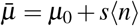:

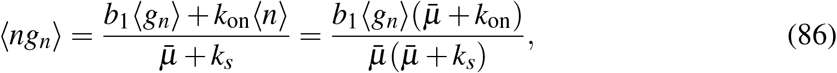

where the second equality uses 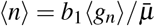 .

### Exact solution

Substituting Eq. (86) into the covariance and the promoter steady-state equation gives a single nonlinear equation in ⟨*g*_*n*_⟩:

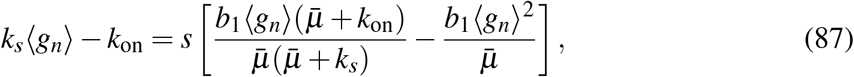

with 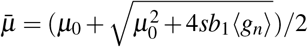. This equation is solved numerically for ⟨*g*_*n*_⟩ at each value of *s*, and the enrichment ratio is

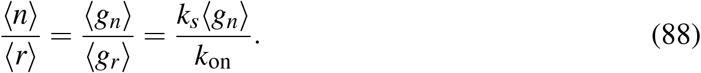

This solution is exact at the level of second moments: no moment closure approximation is needed because the universal cancellation eliminates all terms involving ⟨*n*^2^*g*_*n*_⟩ or higher. The only approximation is the neglect of correlations between cell volume and intracellular state, which is the same continuous-dilution framework used throughout the paper.

Expanding Eq. (87) to first order in *s* recovers the perturbative result (Eq. 50): ⟨*n*⟩*/*⟨*r*⟩ ≈ 1 + *s*(*F*_0_ − 1)*τ*_*g*_, where *F*_0_ = 1 + *b*_1_ *p*_off_*/*(*µ*_0_+*k*_*s*_) is the neutral Fano factor and *τ*_*g*_ = 1*/k*_*s*_. At larger *s*, the exact nonlinear solution captures the saturation of enrichment as the promoter ON fraction ⟨*g*_*n*_⟩ approaches unity, where the perturbative expansion breaks down (Figure 4c).

